# Reproducing within-reef variability in coral dynamics with a metacommunity modelling framework

**DOI:** 10.1101/2024.01.21.576579

**Authors:** Anna K Cresswell, Vanessa Haller-Bull, Manuel Gonzalez-Rivero, James P Gilmour, Yves-Marie Bozec, Diego R Barneche, Barbara Robson, Ken Anthony, Christopher Doropoulos, Chris Roelfsema, Mitchell Lyons, Peter J Mumby, Scott Condie, Veronique Lago, Juan-Carlos Ortiz

## Abstract

Reef systems span spatial scales from 10s to 100s and even 1000s of kilometres, with substantial spatial variability across these scales. Managing and predicting the future of coral reefs requires insights into reef functioning at all spatial scales. However, investigations of reef functioning often consider individual reefs as the smallest unit (10s of kilometres), despite substantial spatiotemporal variability occurring within-reefs (100s of meters). We developed *C∼scape,* a coral metacommunity modelling framework that integrates the demography of corals with population-level responses to physical and environmental spatial layers, to simulate a mosaic of interacting coral communities across a heterogenous seascape. Coral communities are linked using biophysical connectivity modelling. Coral community growth is modelled with a logistic growth model, with the intrinsic growth parameter determined from taxa-specific Integral Projection Models to incorporate demographic mechanisms. Site-specific coral habitat parameters, derived from satellite-based geomorphic and benthic habitat maps, define the maximum coral cover and are used to modulate community growth spatially and temporally as a function of the available space suitable for corals. These parameters are a proxy for the many interacting physical and environmental factors — e.g., depth, light, wave exposure, temperature, and substrate type — that drive within-reef variability in coral demography. Using a case study from the Great Barrier Reef, we show that modulating community growth using site-specific habitat parameters enables more accurate hindcasts of coral cover dynamics, while overlooking within-reef variability may lead to misleading conclusions about metacommunity dynamics. More generally, *C∼scape* provides a valuable framework for predicting spatiotemporal dynamics of coral communities within and between reefs, offering a mechanistic approach to test a range of management and restoration options.

## Introduction

Complex ecosystems such as coral reefs are governed by processes occurring across a range of spatial and temporal scales. Predicting the future of coral reefs to inform management decisions today requires understanding of their functioning across all these scales. A considerable body of literature establishes the importance of variation between reefs, at scales from 10s to 1000s of km, and elucidates how physical and environmental factors influence coral composition, functioning and temporal dynamics [1–7]. Similarly, substantial variation also exists within-reefs, most obviously manifesting as differences between reef zones (i.e. reef slope, flat or lagoon, [8–10]), but also varying locally at finer resolutions, from meters to kilometres. Local physical and environmental factors, such as depth, light, wave exposure, water circulation, temperature, and substrate type drive variability within-reefs [11–14].

Understanding within-reef variability is crucial, as coral populations in different parts of a reef may have varying influence on the overall growth, survival, and recovery of a reef [15]. Assuming one part of a reef is indicative of the entire reef’s functioning, or calculating an average of several sites without accounting for the distribution of coral populations, can lead to inaccurate conclusions about the overall health and performance of the reef [16].

Furthermore, in the context of applied conservation, some locations may provide more optimal restoration or management areas [16–18]. However, obtaining insights into within-reef variability poses challenges. The complexity of processes occurring within-reefs are rarely supported by field data or modelling at a scales that capture dynamics across the heterogenous seascape of a whole reef [19].

With a mechanistic modelling framework, complex processes that occur within-reefs, spanning from individual corals, populations and metapopulations, that are interconnected across seascapes, can be investigated [20, 21]. Several processes are key to such a framework. On a reef, corals are recruiting, growing, reproducing, and dying, through a life cycle that has evolved over millennia. These demographic processes determine population growth rate and can vary spatially and temporally in response to the local environmental conditions [1, 21–24]. Additionally, stress experienced during acute disturbances varies spatially across a reef due to the localised physical and environmental conditions [25–29]. For example, shallower sites with low water circulation often experience greater stress during a marine heatwave [29–31]. Populations that experience greater mortality during disturbance may be repopulated from remnant upstream populations [e.g., 6, 32, 33], and thus it is necessary to consider larval connectivity across metapopulations.

A demographic foundation to a mechanistic modelling framework for coral offers many opportunities for advancing coral reef research [21, 22, 34–36]. Integral Projection Models (IPMs) use empirical information on an individual’s vital rates – survival, growth and reproduction – to estimate population statistics such as growth rate [37, 38]. IPMs have been applied to many species across the animal and plant kingdoms but have mostly been used to consider single populations of a species in one location [39–41]. More recently, the application of IPMs for metapopulations has gained momentum [42, 43]. Metapopulation modelling using IPMs offers potential to mechanistically investigate how changes to individual corals may feed through to a population and metapopulation across space and time.

There are, however, challenges with using projection models for long-term prediction. IPMs were originally designed to explore the population size frequency distribution at equilibrium, and therefore are not intrinsically constrained in their rates of growth, survival or fecundity by limiting factors [22, 44, 45]. For example, a population of sessile organisms, such as corals, cannot have indefinite population growth due to space limitation, and therefore population growth rate needs to be modulated by available space [16, 45–47]. Furthermore, IPMs can require large amounts of demographic data, particularly when trying to represent environmental gradients, although there are methods to circumvent data gaps [37, 48].

Population dynamics are often described with logistic ordinary differential equations, which have a growth rate parameter and are limited by the availability of a critical resource, such as suitable habitat [47]). This differential equation approach has been applied successfully across many marine ecosystems [47, 49–52], but doesn’t capture the underlying demography mechanistically, i.e., how is a particular process in the life cycle, growth of adults or recruitment, for example, influencing the overall population growth? If demographic information on growth, survival and reproduction is available, the population growth rate from an IPM can be used in an ordinary differential equation describing population change with time [46]. Such population models can then be combined into a community model, by accounting for the common utilisation of the limiting resource (space) by the different populations [47]. This approach can be used to create a community model with a more mechanistic foundation to demography, while still capturing the influence of resource depletion as the maximum population size is approached.

The distribution and abundance of corals across a reef is a product of multiple interacting physical and biological processes, which collectively constrain the maximum coral cover and the rate at which that upper threshold is reached [16, 47]. Remotely sourced data that can distinguish variability in environmental, physical and biological parameters offers an option for constraining population growth rate as resources become limited [53, 54]. Geomorphic and benthic habitat maps have been used to estimate the coral habitat on the Great Barrier Reef [55]. These maps can also facilitate partitioning reefs into smaller sites of relatively homogenous physical and environmental conditions. Surrogates, or proxies, have been used to explain the distribution and composition of marine sessile organisms [47, 56, 57]. Given that available substrate within-reefs has been tightly linked to environmental drivers like wave exposure and temperature, modulating coral community growth using spatially explicit available space, or ‘coral habitat’, may act as a proxy for the many processes that determine the distribution of corals and their demographic rates [see 16, 47].

In this study, we built a mechanistic metacommunity modelling framework that integrates the demographic rates of corals, using integral projection models, with community-level responses to spatial and temporal environmental variation via a logistic ordinary differential equation model. We utilise high-resolution habitat maps to construct a spatially explicit seascape of interacting coral populations across a mosaic of sites, each containing two distinct coral types forming a simple community. Across a cluster of reefs, we establish connections among sites to create a ’metacommunity’ using fine-scale larval connectivity modelling. Coral population growth rates from integral projection models inform a logistic growth community model, which adjusts for reductions in growth as the maximum available coral habitat is approached. We compared two parameterisations for coral habitat: assuming a uniform distribution across the reef versus considering spatial variation through habitat maps. For the former, we assumed a maximum coral cover of 80%, recognizing varying assumptions in previous studies regarding this upper limit (e.g. 100% [4, 50], 50-90% [5], or parameterisation from empirical data [16, 58]). For the latter, we developed a methodology deriving a site-specific coral habitat parameter from the habitat maps, , exploring its potential as a proxy for processes influencing coral distribution and population growth across a reef.

We tested both parameterisations in a case study of a cluster of five reefs around Moore Reef in the central/northern Great Barrier Reef. Testing both parameterizations in a case study of a cluster of five reefs around Moore Reef in the central/northern Great Barrier Reef, we assessed whether the site-specific coral habitat parameter captures within-reef variability and allows more accurate hindcasts of long-term reef trajectories than assuming uniform coral habitat across the entire reef. We contextualize how within-reef variability may influence conclusions about average reef dynamics.

## Results

### *C∼scape* model framework

The *C∼scape* modelling framework (Figure 1) combines coral demography with the major processes that vary spatially within-reefs and influence coral population growth, to project coral metapopulation dynamics (Figure 3).

**Figure 1.**
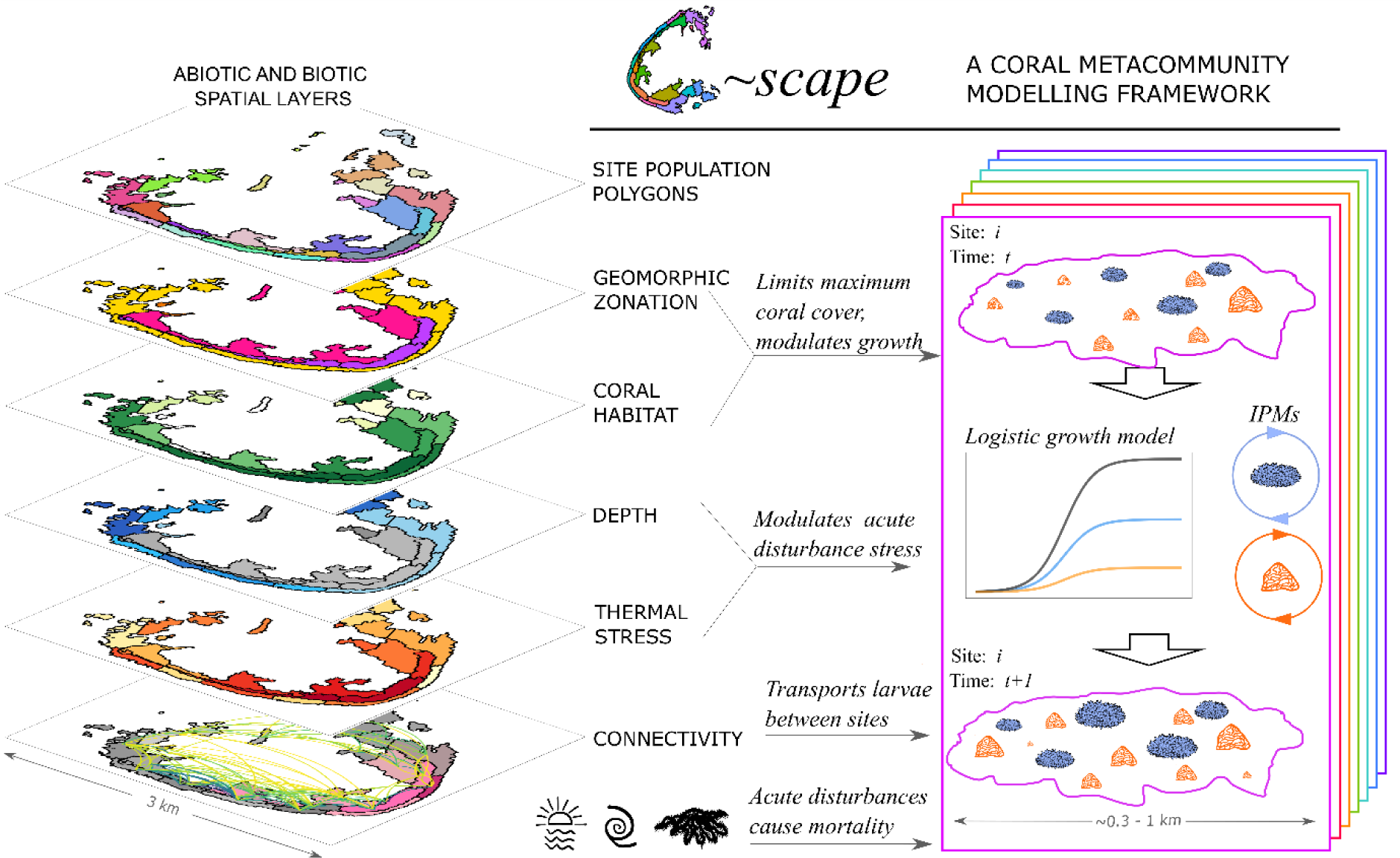
Schematic representation of the C∼scape framework. C∼scape is applied to a reef or cluster of reefs. The reef(s) are divided into spatial units for modelling, referred to as ‘sites’. A coral population for each of two distinct coral types is modelled discretely at each site, with these populations connected with a connectivity matrix representing the probability of coral larvae moving between sites. The abundance and size of corals in each site are tracked. Sites have different depths, influencing the populations’ exposure to temperature stress. Information on thermal stress variability and connectivity between sites is required. Intrinsic population growth rates are determined for each coral type from integral projection models (IPM) that describe the coral life cycle, enabling the projection of changes in each population of corals over an annual period from time t to time t+1. The IPM contains discrete life states for eggs, larvae and settlers, as well as a continuous state across the size range of the coral types. Sites are assigned a value for the ‘coral habitat’, which specifies the maximum percentage total coral cover in a site — this value modulates community growth as the maximum is approached. Acute disturbances — temperature stress, cyclones and crown-of-thorns starfish outbreaks — affect coral populations by killing individuals.

### A spatially explicit seascape

Our methodology for using geomorphic habitat maps to partition reefs into a heterogenous seascape was used successfully on a set of five reefs, the ‘Moore Reef Cluster’, offshore from Cairns on the central/ northern Great Barrier Reef. These reefs were partitioned into 213 sites (Figure 2), each approximately 300-700 m in diameter. A simple coral community (composed of two distinct coral types) was modelled within each site. Long term monitoring data [59] allowed the comparison of the modelled total coral cover trajectories with those from fixed-position transects and from manta tow surveys at model sites overlapping long-term monitoring sites (Figure 2, see more detail in Methods).

**Figure 2.**
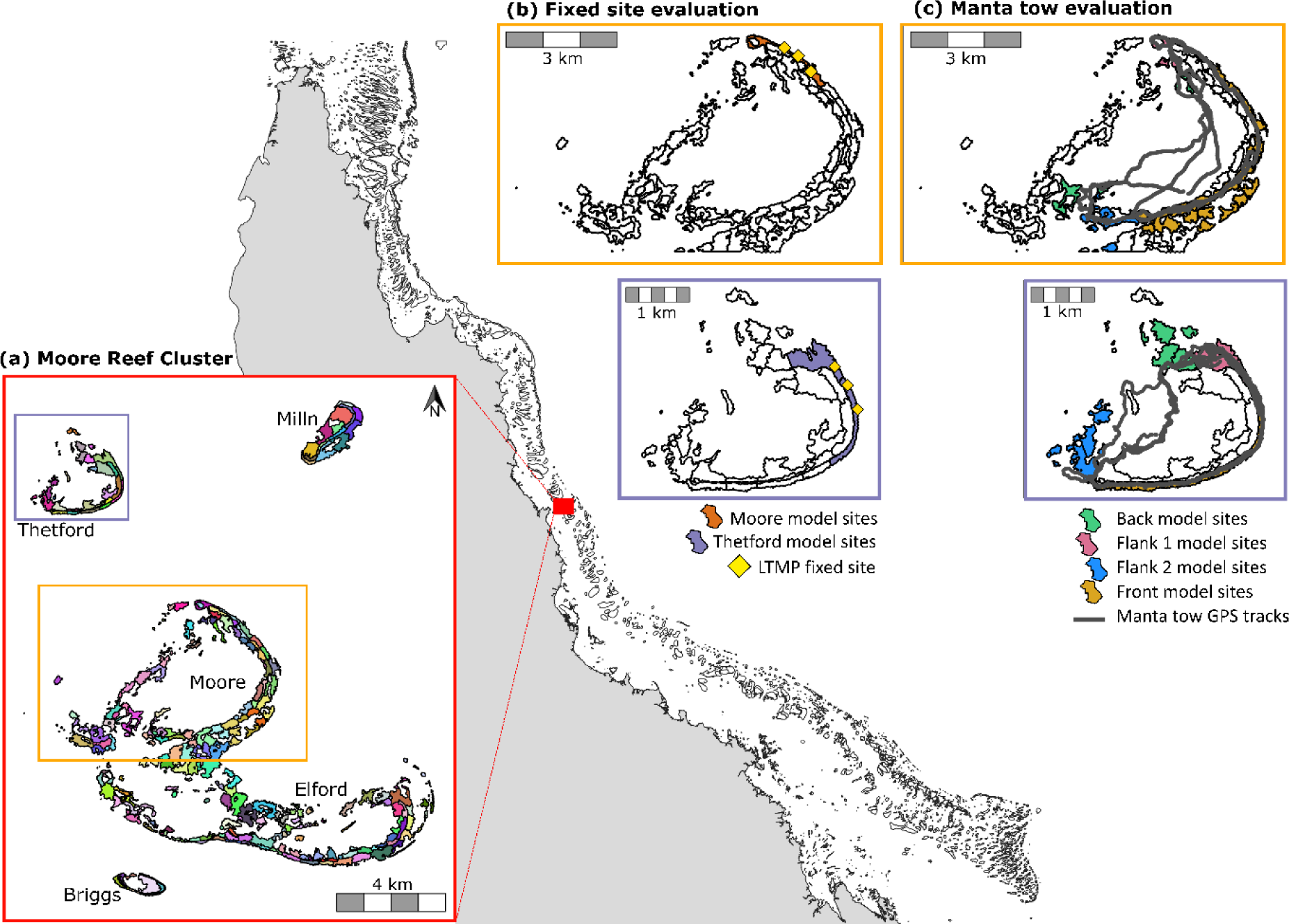
a) The ‘Moore Reef Cluster’ case study showing the reefs broken into sites. Each of the five reefs is labelled, and Moore reef and Thetford reef are delineated by rectangular outlines as they had AIMS long-term monitoring data available for model evaluation. The location of these monitoring locations, and the model sites for which we conducted comparative analyses, are shown in b) and c) for the fixed-position photo transect dataset and the manta tow dataset. In b) yellow points show the location of the three fixed-position transects at each of the two reefs with coral cover data available. In c) the manta tow tracks are shown in grey. The sites underlying these tracks, from which model data were compared to this empirical data for the population growth rate, are one of four colours representing the four manta tow sectors – back reef, flank 1, flank 2 and the reef front – or white, meaning they were modelled but not included in comparison with LTMP data.

**Figure 3.**
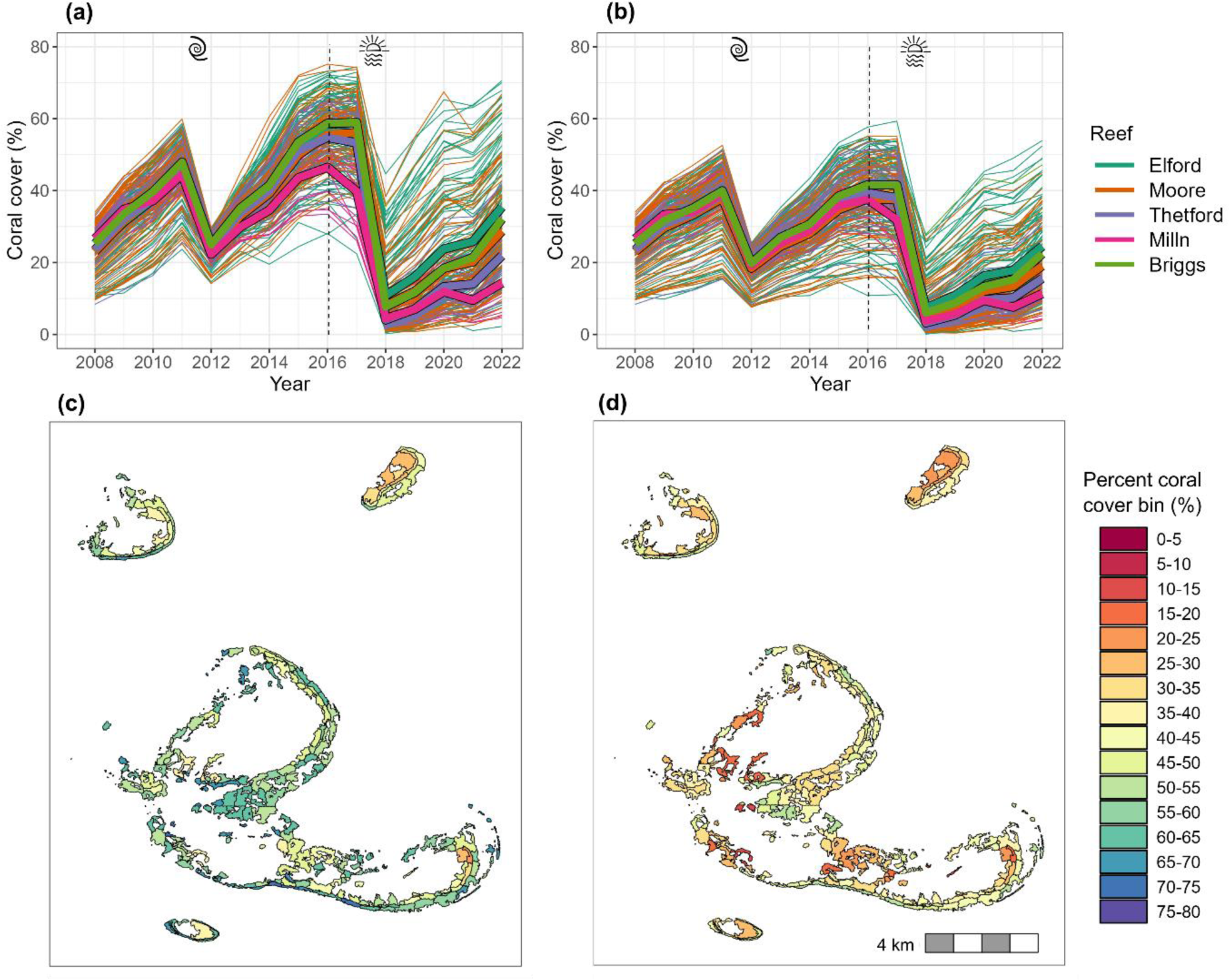
a), b) Trajectories in coral cover between 2008 and 2022 where the model was informed by a) assuming uniform coral habitat availability across the reef cluster and b) site-specific coral habitat derived from benthic habitat maps. Each narrow line represents a different site, with colour indicating one of five reefs in the cluster. Thick coloured lines bordered in black show the site average for each reef. (c) and d) show the spatial patterns in coral cover in 2016 for the uniform and site-specific coral habitat availability scenarios, respectively.

### Spatiotemporal coral dynamics

Applying the *C∼scape* framework to the Moore Reef Cluster allowed us to hindcast dynamic coral cover trajectories. Spatial variation in these dynamics can be attributed to the interactions between population growth, disturbance, fine-scale variation in coral habitat, connectivity and temperature stress. The impacts of acute disturbances were evident through sharp declines in coral cover between 2011-2012 (cyclone) and between 2017-2018 (temperature stress), followed by recovery (Figure 3a, b).

Here we focus on the coral habitat parameterisation for modulating coral population growth within-reefs. In comparing the model hindcasts for the two parameterisations of coral habitat (uniform value of 80% cover for all sites, versus the benthic map derived site-specific values, Figure S2), we found higher maximum coral covers and faster recovery rates (as time- averaged annual change in coral cover) when using the uniform parameterisation, compared to the site-specific parameterisation (Figure 3a vs b).

Variation in coral cover between modelled sites across the reef cluster was evident, with covers diverging by more than 40% between sites when the site-specific coral habitat parameterisation was used, particularly several years following disturbance (Figure 3b). Variation among reefs was also present in both coral habitat parameterisations (with more variation under the uniform parameterisation), but far less than the variation between sites (Figure 3a, b).

Examining the spatial distribution of coral cover across the reef cluster at a point in time (e.g., 2016, Figure 3c, d) shows the spatial variability predicted across the reef by *C∼scape*.

### Reproducing observed coral dynamics

Using the site-specific coral habitat parameterisation tended to improve the match between modelled trajectories and the LTMP observations from both fixed-position photo transects and manta tow surveys (Figure 4a, b). While the overlap in the model predictions and the observed LTMP trajectories was greater when the site-specific coral habitat parameterisation was used, the LTMP trajectories, in particular the manta tow data, had high variability between years, due partly to this being a rapid survey method that does not survey the exact same parts of reef each year, and thus complicating comparisons (Figure 4c, d). Examining the time-averaged annual change in coral cover in the absence of acute disturbances therefore allowed a more quantitative comparison (Figure 5).

**Figure 4.**
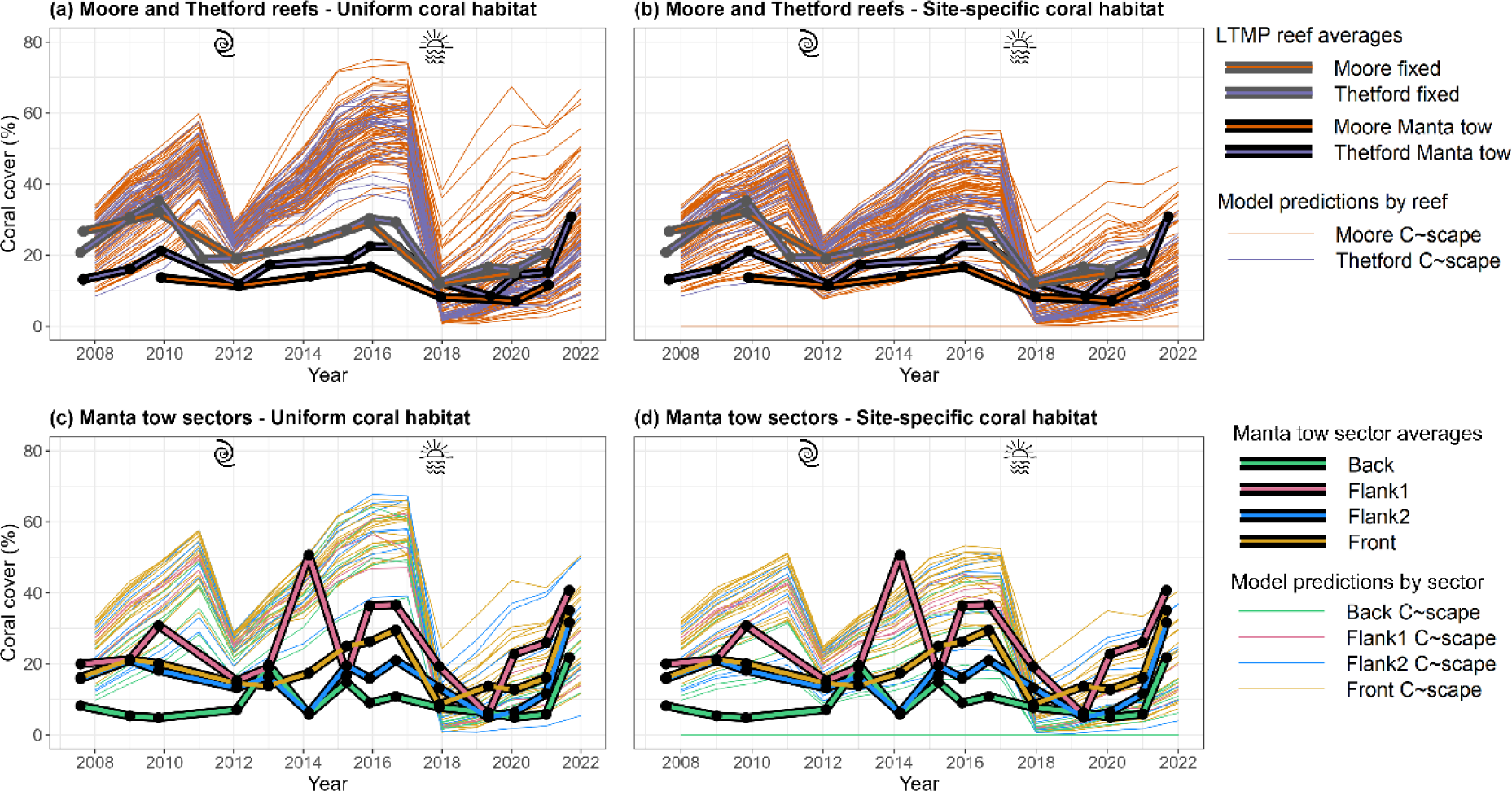
Reef level dynamics for Moore and Thetford reefs (a and b) and manta tow reef sector dynamics within Moore and Thetford reefs (c and d). Narrow lines show the trajectories of different sites in the model with colour indicating reef in a) and b) and sector c) and d). Thick lines in a) and b) show the mean trajectories observed in the LTMP data for the two reefs, with bordering in grey showing the dynamics of the fixed position photo transects and bordering in black showing the mean trajectories of the manta tow data. Thick lines in c) and d) show the mean trajectories of the manta tow data broken into the four sectors and averaged across the two reefs. Variability associated with the empirical data can be viewed in Figure 7. Left panels show the model output under the spatially uniform coral habitat parameterisation, while right panels show the modelled trajectories when coral habitat was parameterised to be site-specific using benthic habitat maps.

**Figure 5.**
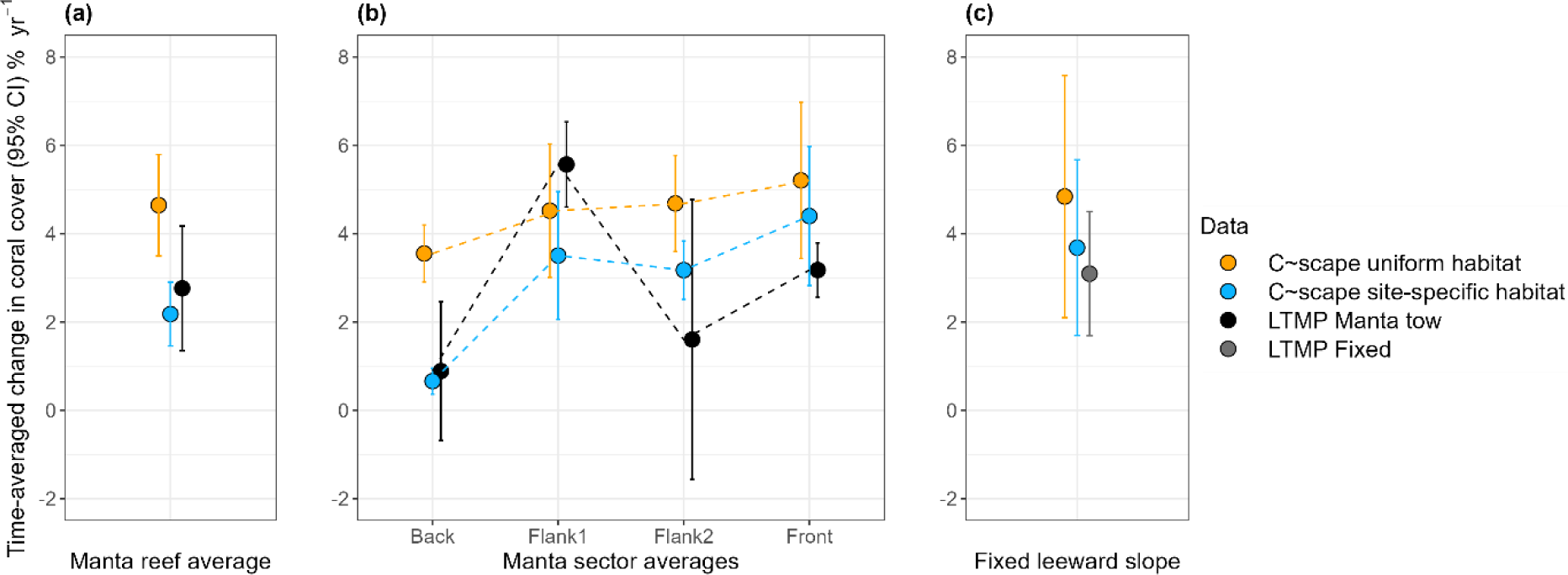
Time-averaged annual change in coral cover in the absence of acute disturbance as percent change in coral cover per year for the growth windows shown in Figure 7. The first panel shows the predicted time- averaged annual change in coral cover for all sites underlying the manta tows, averaged across all growth windows, informed by the two different coral habitat parameterisations (uniform and site-specific), and the average observed time-averaged annual change across all manta tow observations. The second panel shows the time-averaged annual change in coral cover broken down into the four different manta tow sectors. The dotted lines are included to aid visual interpretation of the pattern of variation between sectors across the different datasets. The third panel shows the time-averaged annual change in coral cover at the LTMP fixed sites compared to the predicted time-averaged annual change in coral cover for the six polygons directly underlying these fixed sites at Moore and Thetford reefs.

### Time-averaged annual change in coral cover

According to the manta tow data, the time-averaged annual change in coral cover across Moore and Thetford reefs in the absence of acute disturbance was 2.76±1.41% coral cover yr^-1^. The time-averaged annual change in coral cover predicted by *C∼scape* for sites underlying the manta tows, under the site-specific parameterisation of maximum coral habitat, was approximately 20% slower than of the observed manta tow rate, at 2.24±0.72% yr^-1^ and with the confidence interval overlapping the manta tow observations (Figure 5a). In contrast, under the parameterisation of uniform coral habitat, the time-averaged annual change was more than 1.6 times faster, at 4.49±1.20% yr^-1^ with less overlap in the confidence intervals (Figure 5a).

According to the fixed position transects on the leeward slope, the time-averaged annual change in coral cover was 3.09±1.41% yr^-1^ in the absence of acute disturbance. The model predicted a slightly higher rate at 3.78±2.04% yr^-1^ when parameterised with the site-specific coral habitat values, and higher again when uniform coral habitat was assumed (4.84±2.74% yr^-1^, Figure 5c)

The time-averaged annual change in coral cover for modelled hindcasts using the uniform coral habitat parameterisation showed some variation in rate between the different manta tow sectors of the reef (Figure 5b), with the Front sector having a slightly faster rate than Flank 1 and 2, which in turn had a faster rate than the back sector. These differences are not driven by the site-specific coral habitat parameter, given this is uniform across the reef, and can therefore be assumed to be driven by larval connectivity or variation in post-disturbance coral cover at each site due to temperature stress variability.

Differences between sectors were more pronounced when site-specific coral habitat values were assigned from the benthic habitat maps. In this case, we predicted differences in the time-averaged annual change in coral cover between the manta tow sectors that matched more closely to the empirical data. *C∼scape* predicted fastest recovery rates in the Front and Flank 1 sectors, while Flank 2 had a slightly lower rate and recovery in the Back reef sectors was the lowest. The time-averaged annual change in coral cover at the Back sector matched closely between empirical and modelled data, while *C∼scape* somewhat overestimated recovery rate on Flank 2 and Front sectors but underestimated on Flank 1. When comparing the time-averaged annual change in coral cover for LTMP fixed-position photo transects on the leeward slope of Moore and Thetford reefs, we found that *C∼scape* overestimated the annual change in cover in this location, but less when using the site-specific habitat parameterisation.

*C∼scape* produced outputs at a finer spatial scale than captured in the available empirical data. Therefore, these outputs cannot be validated at the site scale at Moore Reef, but it is still valuable to examine the model predictions of within-reef variability. Variability between sites temporally dynamic and was notably influenced by disturbance. Figure 6 summarises the variation in coral cover in the *C∼scape* output at different times relative to disturbance.

**Figure 6.**
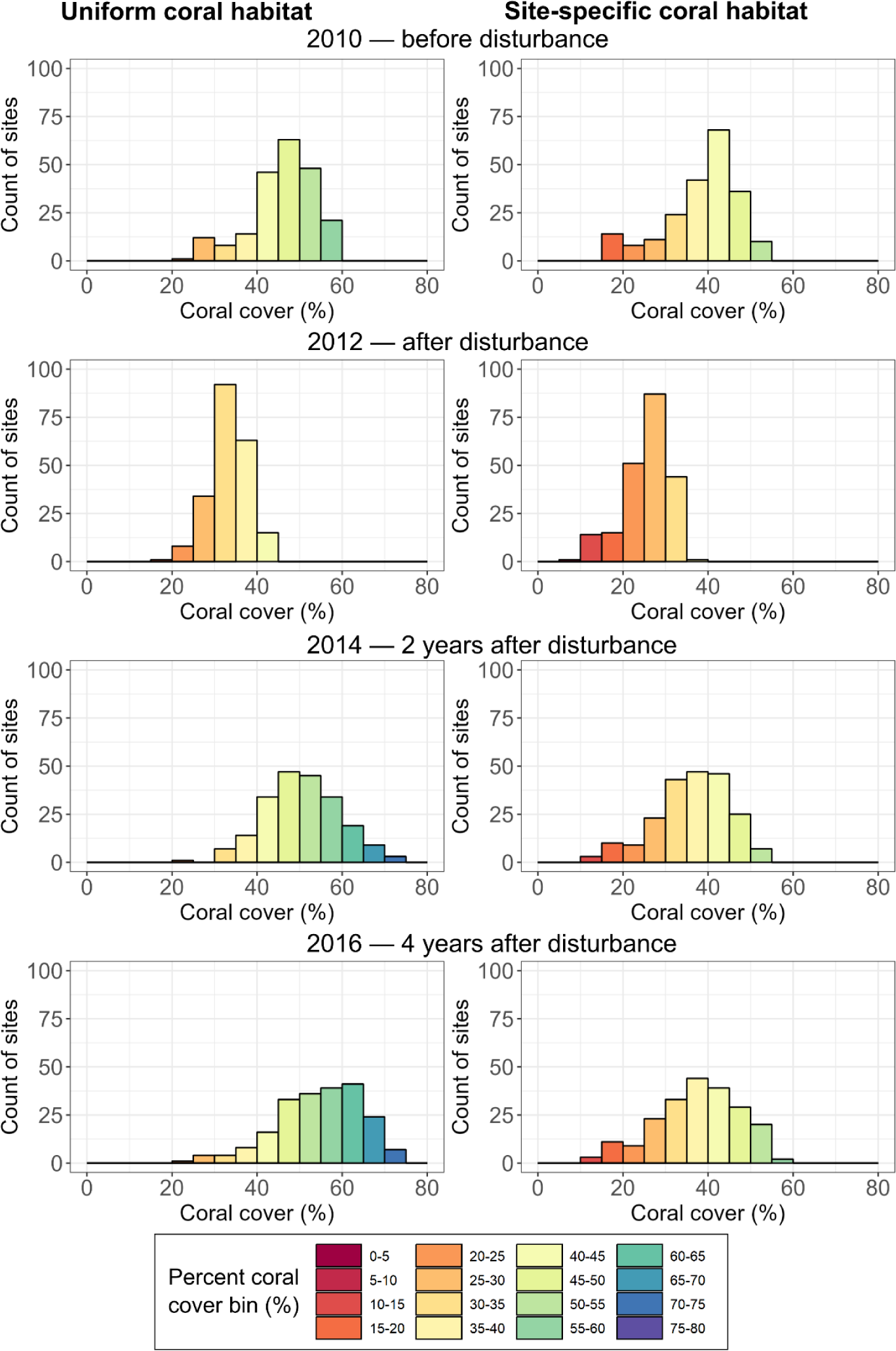
Histograms showing coral cover across sites in the Moore Reef Cluster in four different years: 2010 (prior to disturbance), 2012 (the year after disturbance), 2014 (∼2 years after disturbance), 2016 (∼4 years after disturbance). Left panels show the distribution of coral cover when the coral habitat is assumed to be uniform across all sites with a value of maximum 80% coral cover. Right panels show the distributions when coral habitat was site-specific and informed from the benthic habitat maps.

Disturbances are seen to cause a reduction in the variation in cover and a shift to lower coral covers. The greatest variation within the reef was realised at the longest time since disturbance, ∼4 years following the cyclone event in 2011, when the model was parameterised with the site-specific coral habitat values.

As well as influencing population growth rate and maximum coral covers, the use of the site-specific parameterisations also had implications for the estimated area of coral that could be obtained across the modelled reef cluster. In the Moore Reef Cluster, the total modelled area, i.e., the sum of all site polygon areas, was 2259 hectares. When we assumed uniform coral habitat of 80% across the sites, the total possible live coral area would be 1807 ha, whereas informing the maximum coral habitat for each site based on the benthic habitat maps would give a total area of 1207 ha.

## Discussion

We developed a coral metacommunity modelling framework, *C∼scape*, with the goal of capturing the effects of fine-scale variation in coral habitat, acute disturbances, and larval connectivity between sites in a cluster of reefs. We produced coral community dynamics for 213 discretely modelled sites, linked by larval connectivity across a case study reef cluster containing five reefs. By accounting for fine-scale variation in coral habitat suitability, heat stress, and connectivity, we matched hindcast projections for coral populations within four reef sectors over more than a decade (2008 – 2022), including capturing impacts from a coral bleaching event and a cyclone. The model framework includes many processes, but in this study, we focused on investigating whether incorporating a site-specific parameter for the suitable coral habitat could be used to spatially modulate a general community growth model and improve the ability to hindcast spatiotemporal dynamics. We build the model framework with a mechanistic foundation, structured around demographic processes, by using integral projection models (IPMs) to determine the intrinsic population growth rates for two coral types composing a simple coral community. While the most mechanistic approach to incorporate variation in demographic rates due to environmental gradients would be to include covariates such as depth, temperature, wave exposure and space availability into the statistics which underlie the IPMs [34, 48], the amount of high resolution individual-based data for corals is currently limited. We show that within the framework of a combined IPM- logistic growth community model, habitat maps can be used to parameterise maximum coral cover and act as a proxy for many of the processes structuring within-reef variability in coral demographics, in the absence of more detailed information on the many more specific drivers.

Comparing model performance under parameterisations of uniform coral habitat versus spatially explicit coral habitat revealed several important differences in the hindcast coral dynamics. Modelled coral cover dynamics and time-averaged changes in cover were more comparable to the empirical observations in long-term monitoring data when the site-specific parameterisation was used. As anticipated, the use of site-specific coral habitat parameterisation led to greater variation between sites. This variability was temporally dynamic and was notably influenced by disturbance events, suggesting that some of this variability emerged from the interactions within the model, i.e., between the demographic processes and the spatially explicit processes such as connectivity and temperature stress. For instance, there was generally more variability several years following disturbance than immediately after disturbance under site-specific parameterisation, but in the case of uniform coral habitat, this was less pronounced as all sites approached the set maximum total coral cover.

Coral cover dynamics varied between the five different reefs in the case study, but there was generally greater variability within-reefs than between reefs, offering further support for the importance of considering small scale (100s meters) variability. Similarly, empirical studies have shown that differences in environmental gradients across sites within a reef can drive greater differences in ecological interactions than between reefs with similar environmental exposures [60]. Although site-level predictions of coral dynamics could not be evaluated due to a lack of monitoring data at such a fine spatial scale, we were able to examine the performance of the two coral habitat parameterisations in different sectors of Moore and Thetford reefs. The time-averaged rates of change in coral cover had overlapping confidence intervals in all sectors when parameterised with the site-specific coral habitat values. This shows that even amidst other complex interacting processes in the *C∼scape* framework, the inclusion of a site-specific coral habitat parameter that modulates coral demographics can allow within-reef dynamics to be better distinguished at the sector level. While there may be other ways to parameterise and model this variation more mechanistically [48], such as parameterising spatially heterogeneous coral demographics, this would require substantial amounts of data on coral vital rates across environmental gradients.

A version of the satellite-derived habitat maps that were used to calculate the site-specific coral habitat values are available for coral reefs globally [54, 61–63], as well as for other ecosystems [64–66]. Our findings suggest that if accurate, mapping products can be used to generate a metric and proxy for some of the many processes that vary within-reefs. This greatly increases the feasibility of incorporating within-reef variability into other coral reef modelling efforts, which might otherwise not consider coral habitat suitability explicitly or assume it to be uniform across a reef. The approach to utilising estimates of habitat suitability from habitat maps provides an avenue for better representation of fine scale variability in other ecosystems where the main ecosystem engineers are sessile organisms that compete for a space and resources. In marine environments, this could include kelp or mangrove forests and seagrass meadows, or in terrestrial environments, rainforests, or grasslands. The modelling framework allows for the integration of new demographic and environmental data, and improved modelling of disturbances and connectivity, as they become available, which will improve model accuracy and expand the scope of questions that can be explored.

Predictive models aspire to distil a complex system into a more manageable set of general patterns and processes to enable hindcasts and forecasts rooted in ecological behaviour [67]. Increasing the spatial resolution of such models must be done with care, and two conditions are important: (i) the higher resolution model should accurately replicate observed patterns, and (ii) there should be explicit questions or applications of the higher resolution model that remain unanswerable at a coarser scale. In this study, we have provided comparative analysis with long-term monitoring data to show that the *C∼scape* modelling framework can at least partially satisfy the first condition. There is room to improve individual processes in the model, such as the demographic parameterisations or the connectivity modelling, which will allow improved capacity to reproduce observed reef patterns. We have identified a series of critical questions and applications that are only approachable with predictive models at the within-reef spatial resolution we have targeted, thus satisfying point (ii):

- *Extrapolating spatiotemporal dynamics:* Long-term monitoring data is essential for understanding and managing natural systems, but it is costly and therefore there are limits to the spatial coverage and resolution that can be obtained alongside fine-scale temporal data [68, 69]. This means that it is often necessary to generalise and take one part of a reef that has been sampled as representative of the reef more widely. This approach of reef health monitoring underestimates within-reef variability, and likely contributes to communication challenges between scientists, natural management agencies and stakeholders. Field data could be interpreted alongside high resolution within-reef models to extrapolate how measurements represent other parts of the reef.
- *Ecological questions that cannot be answered at a larger scale:* There are many ecological questions that require the consideration of within-reef variability. For example, identifying fine scale refugia from disturbance is dependent on being able to measure or model exposure to disturbance, alongside the potential for larvae dispersal between different locations within a reef. Similarly, identifying the relative importance and interactions between processes such as demography, connectivity and habitat suitability in structuring coral distributions across a reef requires modelling at a within-reef scale.
- *Fine-scale areas of interest for management*: Although many conservation goals are focused on whole-of-system functionality, there are small areas of natural systems that have particular importance for society and therefore are given special conservation attention. For example, even in the case of the Great Barrier Reef, a reef system 2,300km in length, which clearly requires whole-of-reef management actions to maintain its values [70], maintaining ecosystem functioning in small highly utilised areas – for example sites that tourist operators depend on, sites with high cultural values for Traditional Owners, or areas that are critical for fishing activities – requires more locally targeted approaches [71–73]. Furthermore, zoning for use management, e.g., marine reserves, while often covering large scales in the full extent, requires finer-scale boundaries that often partition reefs into different zones. Mechanistic models that focus on within-reef variability offer opportunities to improve the identification of within-reef zoning boundaries to maximise zoning efficiency.
- *Guided deployment of interventions and restoration:* In response to the overwhelming threat posed by climate change, multiple initiatives worldwide are exploring innovative interventions focused at either reducing the magnitude of acute thermal stress or aiding recovery through seeding or planting natural or genetically adapted corals [36, 74–76]. However, logistical and economic constraints significantly limit the area that can be targeted with these interventions. Consequently, identifying the reef(s) that if chosen for intervention would generate the highest system-level benefit is crucial. Yet even once reefs are chosen, identifying areas within a reef to deploy an intervention with the goal of generating the greatest reef-level benefit is also essential, particularly because many of the proposed interventions are deployed at a site scale rather than reef-wide (median restoration size = 100 m^2^ [18]). This motivates the need for fine-scale modelling that can inform the choice of deployment location, and what characterises a site that will lead to the most benefit for the reef(s) [e.g., 17].

There are various ways the benthic habitat maps or other types of maps describing geomorphology and substrate type could be used to calculate the coral habitat parameter used in this study. We presented the methodology for one approach, which created a significant improvement in the predictive accuracy of the *C∼scape* model. Further work could investigate ways to improve the calculation of suitable coral habitat, particularly as the spatial and taxonomic resolution and accuracy of these maps continues to improve.

## Conclusions

We investigated the relevance and implications of within-reef variability, introducing the *C∼scape* framework as a valuable tool for ecological investigations and coral reef management in the Anthropocene. We sought to fill gaps in existing modelling efforts by incorporating spatial scales and biological and environmental mechanisms crucial for effective management and restoration in the 21^st^ century. We integrated high resolution seascape mapping with integral projection models that mechanistically capture demographic processes in the life cycle via a logistic growth community model. We demonstrated some of the model capabilities, in particular illustrating the practicality of using high resolution habitat maps to investigate and model within-reef variation. , the principle of combining newly available high-resolution spatial information (now accessible for coral reefs worldwide [63]) to downscale mechanistic ecosystem models can extend to other marine and terrestrial ecosystems where sessile organisms compete for a space-related resource. There are a wide range of questions for which the model is relevant, which are beyond the scope of the present study, including predicting of reef futures and investigating the relative importance and uncertainty surrounding the other processes contributing to within-reef variability — larval connectivity, depth, wave exposure, spatial heterogeneity in acute stressors and demography. The mechanistic architecture of *C∼scape* lays a strong foundation to explore these questions and advance our understanding of coral reef dynamics.

## Methods

### *C∼scape* model framework

The *C∼scape* modelling framework (Figure 1) combines coral demography with information on the major processes that vary spatially within-reefs and influence coral population growth, to project coral metacommunity dynamics.

Spatial units for modelling were delineated by creating a spatially explicit mosaic of ‘site’ polygons across a chosen geographic area. When delineating these sites, the aim was to reasonably assume relatively homogenous environmental conditions within each site compared to variations between sites (Table 1). Coral populations are modelled separately within each site but are connected via the transport of coral larvae to and from other sites, thus forming ‘metapopulations’.

**Table 1.**
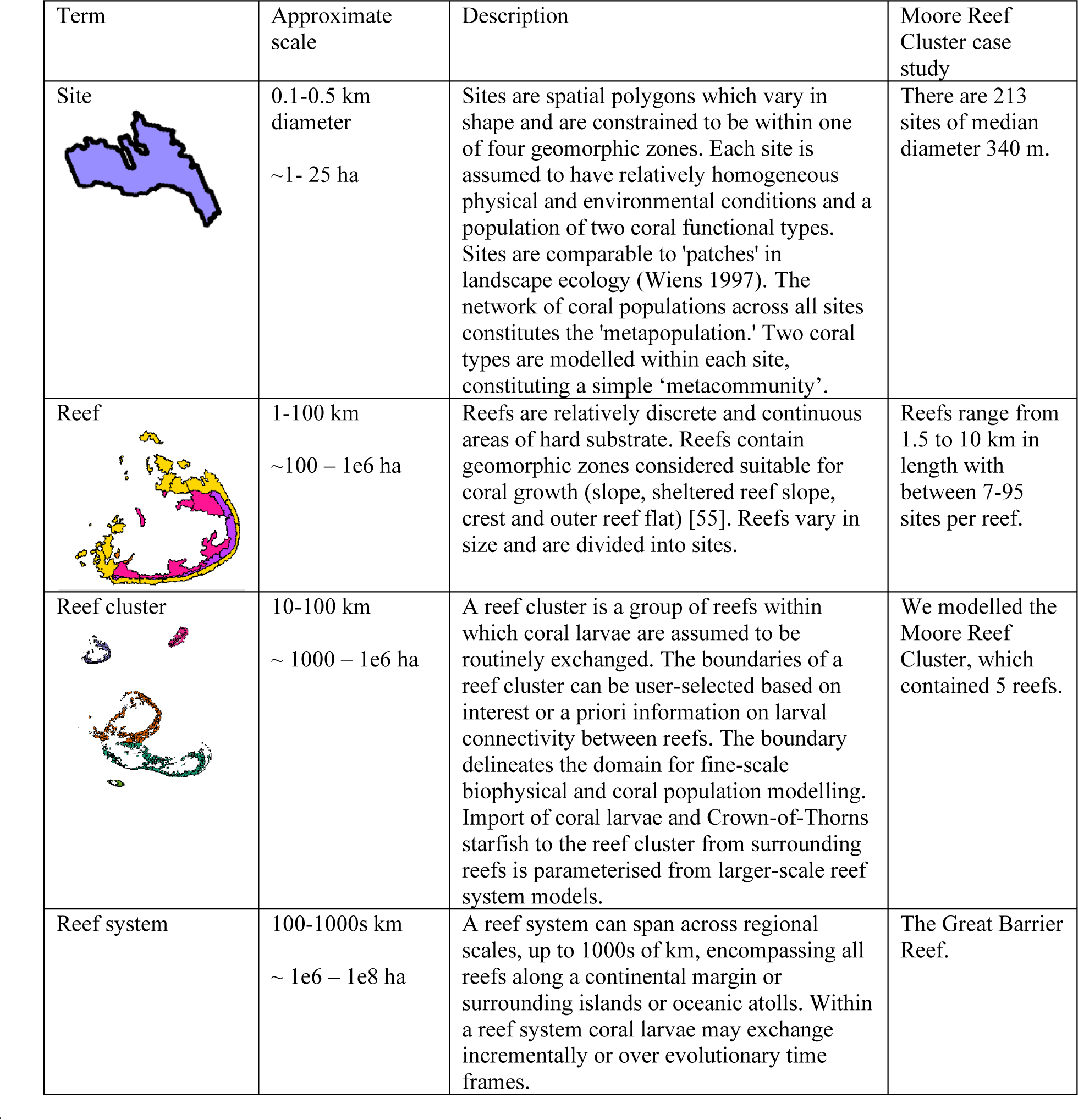
Terminology and description of spatial scales pertaining to the C∼scape model framework, and specifics regarding the Moore Reef Cluster case study.

| Term | Approximate scale | Description | Moore Reef Cluster case study |
| --- | --- | --- | --- |
| Site | 0.1-0.5 km diameter<br>~1- 25 ha | Sites are spatial polygons which vary in shape and are constrained to be within one of four geomorphic zones. Each site is assumed to have relatively homogeneous physical and environmental conditions and a population of two coral functional types. Sites are comparable to 'patches' in landscape ecology (Wiens 1997). The network of coral populations across all sites constitutes the 'metapopulation.' Two coral types are modelled within each site, constituting a simple 'metacommunity'. | There are 213 sites of median diameter 340 m. |
| Reef | 1-100 km<br>~100 – 1e6 ha | Reefs are relatively discrete and continuous areas of hard substrate. Reefs contain geomorphic zones considered suitable for coral growth (slope, sheltered reef slope, crest and outer reef flat) [55]. Reefs vary in size and are divided into sites. | Reefs range from 1.5 to 10 km in length with between 7-95 sites per reef. |
| Reef cluster | 10-100 km<br>~ 1000 – 1e6 ha | A reef cluster is a group of reefs within which coral larvae are assumed to be routinely exchanged. The boundaries of a reef cluster can be user-selected based on interest or a priori information on larval connectivity between reefs. The boundary delineates the domain for fine-scale biophysical and coral population modelling. Import of coral larvae and Crown-of-Thorns starfish to the reef cluster from surrounding reefs is parameterised from larger-scale reef system models. | We modelled the Moore Reef Cluster, which contained 5 reefs. |
| Reef system | 100-1000s km<br>~ 1e6 – 1e8 ha | A reef system can span across regional scales, up to 1000s of km, encompassing all reefs along a continental margin or surrounding islands or oceanic atolls. Within a reef system coral larvae may exchange incrementally or over evolutionary time frames. | The Great Barrier Reef. |

Each population is modelled using a logistic ordinary differential equation modified such that the intrinsic growth rate is substituted for the growth rate obtained from Integral Projection Models (IPMs, [38]. The IPMs are based on the coral life cycle, one for each of two distinct coral types (see Figure S6) and allow predictions of the annual probability of growth and survival, and the amount of reproductive material produced by a population of corals in a site. To account for changes in coral growth and survival due to limits on available suitable habitat, the IPMs are modulated by a coral habitat parameter that defines the maximum total coral cover in a site (further detail in subsequent sections).

Annual environmental forcings that cause coral mortality, i.e., temperature stress, Crown-of-Thorns starfish outbreaks and cyclones are assigned to each site.

### Site population polygons: spatial units for modelling

The geographic boundaries of reefs were determined from a publicly available [77] geomorphic zonation map that depicts the distribution of geomorphic features such as reef crest, slopes, reef flats and lagoons [55]. The geomorphic zonation map extends to a depth of approximately 20 m, and this sets the depth limit to our current modelling efforts.

It is well established that different geomorphic zones are formed by distinctive environmental conditions [15]. Consequently, coral community compositions and dynamics are generally more similar within geomorphic zones than between them [78]. Following the recommendations of Kennedy, Roelfsema (54), we only considered the zones that are expected to have predominantly hard substrate — Reef Slope, Reef Crest, Outer Reef Flat and Sheltered Reef Slope — to be areas where we would expect appropriate habitat and conditions for corals to grow [description in table S1, 55].

Geomorphic zones were further partitioned to account for other spatially varying processes (e.g., connectivity, temperature stress). We pixelated each geomorphic zone into hexagonal cells in R using the geospatial indexing system H3 in package h3 [79] at resolution 12 (approximately 20 m diameter, Figure S1). A distance matrix based on Euclidean distances was then created and the cells were clustered together using the base R function ‘hclust’ with method set as ‘complete’. In this way, hexagons within the same geomorphic zone and adjacent to each other were clustered together to form a polygon that had a maximum longest diameter of ∼1000 m. Polygons did not need to be continuous (i.e., they could be composed of multiple smaller polygons) provided they were in the same geomorphic zone.

### A community growth model using an IPM for intrinsic growth rate

To project coral community growth over annual time steps we used a logistic ordinary differential growth equation as the foundation to the model, which can be written as

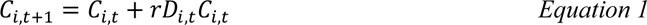

Where *C*_*i,t*+1_ is the total coral cover at time *t+1* at site *i*, *C*_*i,t*_ is the total coral cover at time *t* at site *i*, *D*_*i,t*_ is a parameter accounting for density dependent changes to community growth as the maximum potential coral cover is approached, and *r* is the intrinsic rate of community growth.

We considered populations of two distinct coral types, *ft,* to compose a simple coral community. We wanted to build a mechanistic foundation to the demographic processes captured by the model for these two coral types, and so coral cover was determined from the coral population size structure, 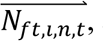, at each time step, for each site:

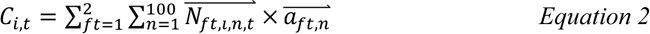

Where *a*_*ft,n*_ is the colony area corresponding to the *n*th of 100 size classes for coral type *ft,* and 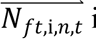 is a vector describing is the abundance of colonies in each size class *n,* at site *i*, at time *t*.

To describe the intrinsic rate of population growth, *r*_*ft*_, for coral type *ft*, we modified the approach in Miller, Jensen (46) and used an integral projection matrix (more detail in following sections) to describe the probability of transitioning between size classes, such that

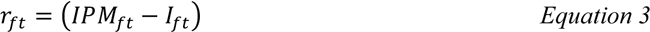

where *IPM*_*ft*_is the integral projection matrix, and *i*_*ft*_ is the identity matrix (see Miller et al. 2002) of coral type *ft*. By subtracting the identity matrix, which is the diagonal of the IPM matrix that describes the probability of a coral colony staying the same size, what remains is the probabilities of growth, shrinkage and survival, and therefore describes what individuals are added/subtracted from the population.

To account for the effect of space limitation on coral community growth, density dependence, *D*_*i,t*_ for site *i* at time *t*, was calculated as:

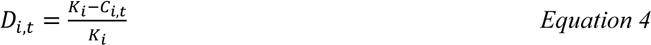

where *K*_*i*_ is the coral habitat (maximum obtainable coral cover) at site *i*, and *C*_*i,t*_ is the total coral cover at site *i* at time *t. K*_*i*_ is designed to act as a proxy for the many interacting physical and environmental factors — e.g., depth, light, wave exposure, temperature, and substrate type — that set the limit on the maximum amount of coral cover that may be obtained.

The final equation to project changes in each coral population *ft*, in terms of size structure, while accounting for the constraints on coral habitat *K*_*i*_, at site *i* at time *t,* is a discrete time- scalar logistic equation:

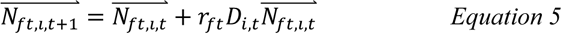

Where 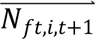 is a vector describing in the number of colonies in each of the 100 size classes at *t+1* and three discrete life stages (eggs, larvae and settlers) ,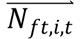 is a vector describing the same at time *t*.

The intrinsic growth rate of each population is multiplied by the community crowding function, such that the processes of growth and survival are reduced as the two populations fill the maximum available coral habitat and the populations will not exceed the maximum coral habitat in a site, *K*_*i*_.

### Integral projection models

The integral projection transition matrix, or kernel *k*(*y, x*), is analogous to a projection matrix (e.g. Leslie matrix, [80]) in matrix population modelling [23]. It represents all possible transitions from state *x* (in year *t*) to state *y* (in year *t* + 1), integrated over all states of *x* [81]:

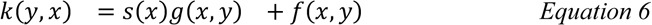

where *s*(*x*) is survival of individuals in state *x* from time t to t+1, *g*(*x*, *y*) is the growth of individuals from state *x* to state *y*, *f*(*x, y*) is the fecundity of individuals in state *x* producing those in state *y* at t+1 [34].

We modelled three discrete states for corals (eggs, larvae and settlers), and a continuous state (coral size) as we wanted to build a mechanistic framework from which the importance of transitions between these discrete states could be evaluated, even if the data available to parameterise some of these transitions are currently limited. Therefore, as shown in Figure S6, we model, or parameterise, the (i) growth and (ii) survival, (iii) fecundity, (iv) fertilisation (the number of eggs that are fertilised and become larvae), (vi) the survival and settlement probability of the larvae (handled separately to the IPM so that (v) connectivity modelling can be used to simulate the exchange of larvae among sites), and (vii) survival of settlers and growth to a 1-year old coral at which point they enter the continuous state (Figure S6, Table S1).

We fit models for each of growth, survival, and fecundity as a function of coral colony surface area (cm^2^) in a Bayesian framework following the approach of Elderd and Miller (82) and Kayal, Lenihan (83) using the brms package in R [84].

Colony growth was modelled by predicting coral colony planar area at time t+1 as a function of coral colony area at time t. Colony survival was modelled by predicting the binary output of alive or dead at time t+1 as a function of colony area at time t. For fecundity, we modelled the number of eggs produced by a coral colony as a function of colony area. We also fitted a model for the number of polyps per cm^2^ coral area as a function of coral size in cm^2^.

Multiplying predictions from these models was used to estimate the number of eggs as a function of colony size (more detail available in S1.1).

Modifying code provided in Kayal, Lenihan (83), we predicted from the regressions for growth, survival and fecundity for 100 coral size classes (spanning from 1 cm diameter to 90% of the largest coral size in the dataset) for each coral type. We converted these predictions into probabilities which formed the integral projection matrices (Supplementary Material 2). The IPM matrices describe the probability of transitioning from and to all 100 size classes and the three discrete life stages (Supplementary Material 2).

### Coral types and data

IPMs require substantial amounts of data to describe the vital rates – growth, survival and fecundity – as a function of a continuous state (here, coral size) with robust statistical regressions. Similar to previous studies [85], we included two distinct coral functional types in the model: ‘corymbose *Acropora’* and ‘sub-massive *Goniastrea’*. While additional coral types would provide deeper insights into community dynamics, these two groups represent two major functional groups on the Great Barrier Reef with different traits: corymbose *Acropora* is a relatively fast-growing coral with lower annual background survival, and relatively high vulnerability to thermal and hydrodynamic stress; in contrast, sub-massive *Goniastrea* are typically slower growing, have high annual survival, and are more resistant to thermal and hydrodynamic stress.

Ideally data for these coral types would come from the location where *C∼scape* is to be applied, but here our focus was to develop a robust framework and so we used the largest datasets available to us. To inform the growth and survival components of the IPMs for corymbose *Acropora*, datasets were utilised from Scott Reef [northwest Australia, 86], Heron Island [Great Barrier Reef, 87], and Moorea, [French Polynesia, 83]. These datasets employed comparable data collection methods, involving the tagging, photographing with a scale, and bi-annual or annual monitoring of tagged coral colonies. Data for sub-massive *Goniastrea* was obtained from the same Scott Reef dataset as for corymbose *Acropora*.

Colony diameter was consistently measured across the three datasets, so we used diameter and converted this to colony area by assuming the colonies were circular.

Information on the number of eggs per coral polyp and number of polyps per colony surface area were available for *Acropora millepora* in the Scott Reef dataset. Additional parameters describing the probability of egg fertilisation, larval settlement, and post-settlement survival for the first year were sourced from the literature (see Table S1).

### Spatially explicit habitat parameterisation

A deterministic IPM assumes that vital rates remain constant over time, which we know is not the case, particularly over extended temporal periods [45]. Therefore, we required capacity to modulate the intrinsic growth rate of the IPM in response to a change in the state of a population and the conditions at a site. Given corals are sessile organisms, space is a limiting resource (Connell, 1961) known to affect population growth rate. To simulate resource limitation as competition for space, the IPM is multiplied by the density dependent parameter, *D*_*i,t*_ (Equation 4), which accounts for the maximum coral habitat at a site. Coral habitat is defined as the maximum percentage cover of corals that could exist across a given site polygon (a comparable approach to that taken by Boschetti, Babcock (16)). In this way the coral habitat acts as a proxy for many interacting factors that influence the habitat suitability for corals.

The maximum coral habitat, as a percentage of the 3D spatial area of each site, is calculated using satellite-derived benthic habitat maps [55] (Supplementary Material 1.2). Each 10 m pixel in the benthic map is classified as one of four categories: Sand, Rubble, Rock or Coral/Algae (see Supplementary Material 1.2).

According to Roelfsema, Lyons (55), pixels defined as dominated by Coral/Algae and Rock can be considered as potential coral habitat with Coral/Algae being “any hardbottom area supporting living coral and/or algae” and Rock being “any exposed area of hard bare substrate”. As such, pixels classified to be dominant in either Coral/Algae or Rock were counted as suitable coral habitat. However, pixels are 10 x 10 m in size, and, due to the patchy nature of reefs at very fine scales, it is unlikely that the full area of the pixel is completely covered by the dominant classification. Hence, we utilised a replicate sampling strategy to account for the uncertainty of within pixel variation. For one of the four possible categories to be dominant, the dominant category would need to cover 26-100% of the pixel, while the non-dominant categories were ≤25%. Therefore, we replicated the calculation of coral habitat 100 times, each time assigning a random value between 26-100% for the dominant category and 0-25% for the non-dominant categories using a uniform distribution. The mean coral habitat from this sampling was calculated as the proportion of potential coral habitat as a percentage. A three-dimensional coral area was determined for each site to consider sloping bathymetry which creates larger substrate area for corals relative to the two-dimensional area (Supplementary Material 1.3).

### Connectivity

#### Within-cluster connectivity

The workflow creates a spatial mosaic of site polygons, and each site represents a distinct area where a population of two types is simulated. To establish a metacommunity comprising these populations, it was necessary to estimate the probability of coral larvae dispersing from one site to another each year. These probabilities are influenced by factors such as physical connectivity, including ocean currents and hydrodynamics during coral spawning, as well as trait and behavioural characteristics of the coral larvae. Such probabilities could be obtained with various modelling or assumptions, but in this case, the eReefs RECOM model [88, 89] nested within GBR1 [90] was used to simulate transport and dispersal of larvae with a ∼250 m spatial resolution.

RECOM is a tool to nest local-scale models within regional-scale hydrodynamic-biogeochemical models of the GBR developed for the eReefs project [89]. RECOM produces a package of initialisation and boundary forcing files extracted from the eReefs environmental models, along with hydrodynamic transport files that can be used to drive more customised model runs. Coral larvae dispersal was simulated using passive tracers released at times that reflected the timing of annual mass spawning events for *Acropora* and *Goniastrea* spp., which typically spawn at the same time of year. Tracers were released on days 4, 5 and 6 following the full moon associated with known spawning events in 2015, 2016 and 2017. Releases were set to occur between 7:30 pm and 11 pm on each night of spawning from the centroid of each site polygon. Tracers were released at the water surface and were assumed to be neutrally buoyant. The ‘competency period’ of larvae was set to be between 4 and 28 days from release for both corymbose *Acropora* and sub-massive *Goniastrea* [91, 92]. The tracers were labelled according to their site of origin and tracked as they were transported by tides and currents through the local reef cluster region from release until the end of the competency period. A zero-gradient boundary was applied to tracer concentrations at the edge of the model domain (i.e., it was assumed that the concentration coming inwards across the boundary was the same as the concentration just inside the boundary).

We made various assumptions in a post-processing workflow to simulate what we expected the larvae would do. We assumed a larvae would settle in the first site it passed over during its competency period. At each hourly time step during this period, the total supply of each tracer was reduced by the proportion of tracer that was over a reef, and the cumulative quantity that had settled on each polygon was tracked. The total amount of tracers settling into each site was then converted into a probability matrix, one for each day of release and for each year simulated. Some tracers were lost from the reef cluster due to advection outside the modelled spatial domain. In the connectivity matrix, this was indicated by columns that did not sum to 1 and we assumed these larvae died or settled on reefs outside our focus cluster.

For the simulations explored in the present study we took the mean of the connectivity matrices for days and for years and used this mean matrix for all years.

#### External larvae supply

It may be reasonable to assume that a cluster of reefs being modelled is a closed system, but this depends on the spatial context. In our case study, the Great Barrier Reef is a well-connected system [5, 6, 93, 94] and there are generally other reefs surrounding potential reef clusters of interest, which likely contribute coral larvae to the reefs inside a chosen cluster. Therefore, for the simulations presented here, the ReefMod-GBR model [5] was also run, for the whole of the Great Barrier Reef, with the same environmental forcings. From the ReefMod-GBR model outputs we could then extract the number of larvae arriving to each reef in the Moore Reef Cluster in each of the years we were interested according to the ReefMod-GBR model outputs.

External larvae were distributed to the sites proportionally based on the total site area, i.e., larger polygons received more larvae. As for larvae arising from within the cluster, these external larvae settled on the reef with a probability dependent on their coral type (Table S1), and then were governed, as for the other individuals, by the IPM and the conditions at the site they were assigned to.

### Acute disturbances

Temperature stress, cyclones, and Acanthaster Crown-of-Thorns (COTS) starfish outbreaks are included as acute disturbances in the *C∼scape* framework. In any given year these disturbances can impact the coral population by killing coral colonies within a site.

#### Temperature stress

Marine heatwaves cause temperature stress and coral bleaching and subsequent mortality [95]. Bleaching mortality can be predicted from Degree Heating Weeks (DHW), a metric of the accumulated heat stress [96]. To create a hindcast of Degree Heating Weeks (DHW) for each site, we first extracted the annual maximum DHW, available at a 5-km resolution from the NOAA Coral Reef Watch (CRW) Product Suite version 3.1 (Liu et al. 2017). Each of the 5 reefs in our case study cluster was assigned a DHW for each year as the maximum DHW value of the nearest 5-km pixel. We then used eReefs RECOM to model the variability in exposure to heat stress across a reef during marine heatwave years (in this case from 1 November in the preceding year through to 30 April in the named year for 2016, 2017, and 2020) (Supplementary Material 3.1). We used only surface DHW, as depth was handled separately in the coral mortality functions.

Taking the DHW assigned to each year and each reef, we then ‘spread’ this value across the different sites within each reef (see Supplementary Material 3.1). We calculated the proportional residuals between the mean DHW of the whole reef and the DHW at each site for the marine heatwave years (Supplementary Material 3.1, Figure S10).

The probability of coral mortality in any given year was estimated as a function of DHW following the relationships developed in Bozec, Hock (5) based on observations of coral mortality during the 2016 bleaching event on the Great Barrier Reef [97]. Mortality is a function of the coral type, the DHW at a site and the depth of a site (see detail Supplementary Material 3.2).

The site polygons were overlaid on a bathymetric map of pixel size 10 x 10 m (https://www.eomap.com/) to determine the mean depth within each polygon, and all corals within each site were assumed to be at this mean depth (Figure S2), which influences the temperature stress they experience (Supplementary Material 3.1).

#### Cyclones

*C∼scape* is set up to receive a storm intensity for each reef defined on the Saffir-Simpson scale (1–5; [98, 99]. Reef level exposure to cyclones can be difficult to determine as it depends on many factors including the distance of a reef from the cyclone path, the speed of the cyclone and fetch distances across which damaging waves can be generated. For this study we therefore assigned a cyclone category based on the observations in the AIMS LTMP which recorded cyclone damage at Moore and Thetford reefs in the year 2011 only. This was caused by Cyclone Yasi, and all reefs in the cluster were assigned a category 2 cyclone in this year, representing the category of Yasi when it passed over the modelled cluster [100].

We did not model variation in exposure to cyclones at the within-reef scale in the present study, nor consider how corals of different sizes may be influenced, but flag these as important areas for future model development.

To model coral mortality due to cyclones we used data from Fabricius, De’Ath (101). We fitted two regressions to the data in fig. 6 Fabricius et al. 2008, one for branching corals, and one for sub-massive corals, to obtain coral mortality as a function of wind speed in m/s (Figure S11). Windspeed was determined from cyclone category according to the Australian Bureau of Meteorology (Supplementary Material 3.2).

#### Crown-of-thorns

COTS population dynamics are not modelled explicitly in *C∼scape*, so instead we received outputs from the ReefMod-GBR model [5] on the predicted COTS density per age class for each reef of the modelled cluster. There were eight age classes with the abundance in each class given as density per *m*^2^. The density for each reef in each year was multiplied by the total site area, i.e., larger polygons received more COTS.

The function for coral mortality caused by COTS follows Bozec, Hock (5) (see Supplementary Material 3.3). We model coral mortality as a function of the abundance of COTS and age-dependent consumption rates. We incorporated a difference in the preference of COTS for the two different coral types, again following Bozec, Hock (5) based on information in De’ath and Moran (102). According to this data, corymbose *Acropora* is preferred over *Goniastrea* at an odd ratio of 14:4.3. Corals of all sizes experienced the same mortality, i.e., a proportion of the total coral colonies was removed by the COTS.

### Case study hindcast

To demonstrate the workflow for applying the *C∼scape* framework to a reef cluster we chose a set of five reefs – Moore, Thetford, Milln, Briggs and Elford – on the Great Barrier Reef, offshore from Cairns (-16.867 °, 146.233 °, Figure 2).

#### Validation data

To determine whether the model framework could capture observed differences in total coral cover trajectories among sites, model predictions were compared to observed time-series of coral cover from the AIMS long-term monitoring program [59, 103, 104] available at two of the five reefs in the Moore Reef Cluster case study: Moore and Thetford reefs. Two datasets were available for these reefs: time-series of total coral cover from fixed-position photo transect data at three sites at on the leeward north-east slope of the reefs, and total coral cover estimates from manta tow surveys spanning the reef perimeters. Manta tow is a monitoring technique where an observer (on snorkel) is towed behind a boat and makes visual assessments, recording the average total coral cover in ∼10 m wide bands on consecutive tows which are ∼200 m in length [104]. The surveys follow the reef perimeter, usually on the reef slope [104]. The manta tow surveys are less accurate than the fixed-position photo transects but provide broader spatial information across the reef. Although designed to give reef level assessment, the manta tow data can be divided spatially into four different reef sectors — back reef, flank 1, flank 2 and front (Figure 2c, Miller et al. 2018) — which was useful for evaluating the within-reef variability produced by *C∼scape*.

#### Model simulations

For this case study we included inputs from the regional-scale model ReefMod-GBR [5] to obtain predictions of external coral larvae and COTS densities in the modelled reef cluster.

ReefMod-GBR outputs were available from 2008 onwards, so we therefore initialised *C∼scape* in this year to examine temporal dynamics through to 2022.

Simulations were initialised based on coral cover recorded in the AIMS LTMP observations from the starting year of the hindcast (2008 for the full trajectory, or the start of each ‘recovery window’ (Figure 7) for the population growth rate analysis, see S2.1).

**Figure 7.**
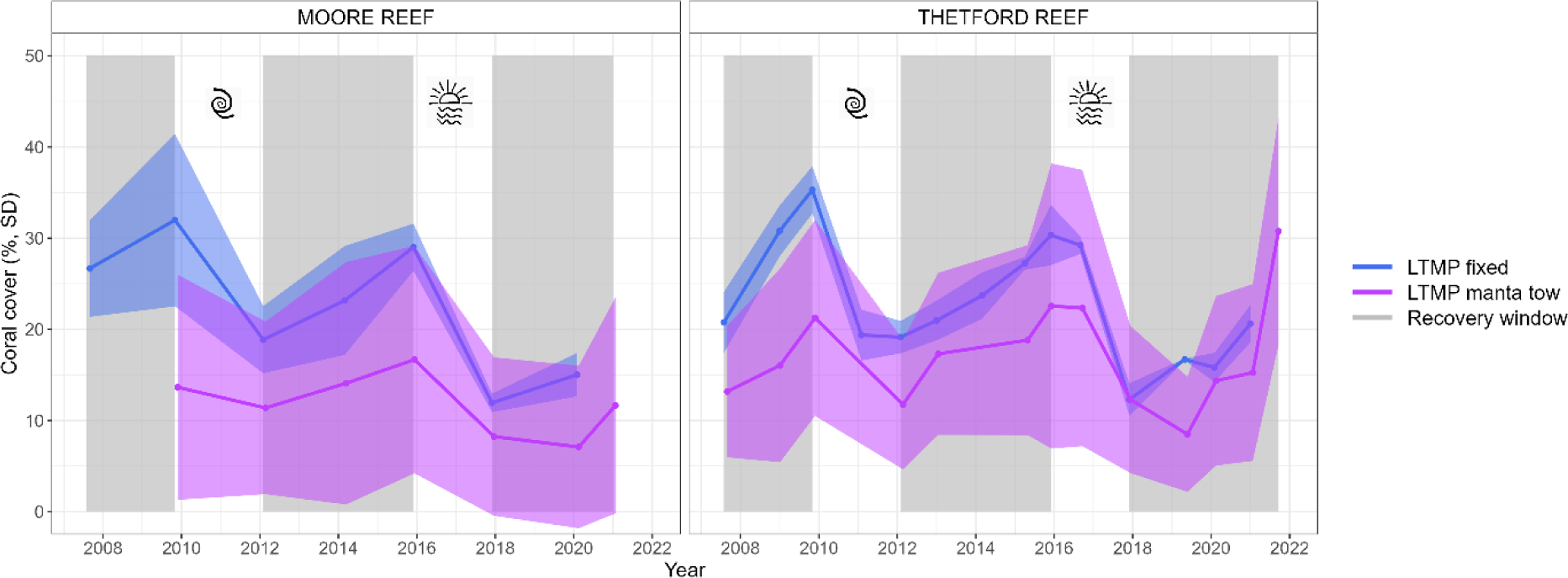
Temporal dynamics of the AIMS LTMP fixed-position photo transect and manta tow data in blue and purple, respectively. Mean and standard deviation are calculated from the three sites in the LTMP fixed-position photo transect data and across all available tows for the manta tow data. Two major disturbances occur in the time-series, a cyclone in 2011 and a marine heatwave in 2017, as shown by symbols. Grey shading shows the ‘recovery windows’, i.e., time-periods identified to investigate population growth rate in the absence of acute disturbance.

We ran the model for two parameterisations of maximum coral habitat for each site in the reef cluster: uniform and site-specific coral habitat. For the uniform parameterisation of coral habitat, we tested a scenario where the maximum coral habitat across was set at 80% for all sites. Various assumptions have been made regarding this upper limit in previous modelling studies (e.g. 100% [4, 50], 50-90% [5], or it is sometimes informed from empirical field data [16, 58]. For the second scenario, we developed a methodology for calculating a site-specific coral habitat parameter from satellite derived geomorphic and benthic maps. In this way we could investigate whether a site-specific parameter describing the availability of coral habitat could act as a useful proxy for the many processes that ultimately determine coral distribution and population growth. We wanted to contrast how using the available benthic habitats may improve upon a basic assumption of the maximum coral habitat.

We examined the full trajectory of the modelled hindcasts (for uniform coral habitat and for site-specific coral habitat calculated from the benthic maps) for all sites in each of Thetford and Moore reefs. We compared these trajectories to those observed in the manta tow and fixed-position photo transect time-series.

The manta tows were classified as one of four sectors. Therefore, model sites were also classified as one of these four sectors: any polygons spatially underlying the manta tow tracks based on GPS tracks of the manta tow were assigned to the underlying manta tow sector, provided they were also classified as the reef slope or sheltered reef slope geomorphic zone as the manta tows do not survey the outer flat or the crest habitats (Figure 2b,c). We visually compared trajectories of total coral cover in the empirical manta tow data for each sector with the predictions of total coral cover for the underlying *C∼scape* polygons.

All scenarios were replicated 100 times, each sampling once from the 100 the integral projection matrices generated to capture demographic uncertainty.

#### Time-averaged annual change in coral cover

Validating time-series dynamics can be challenging because, for instance, if a modelled disturbance early in the time-series leads to a perturbation that is either smaller or larger than observed in the empirical data, it introduces errors that propagate throughout the analysis. (Bozec et al. 2022). Therefore, we identified temporal windows without acute disturbances and reinitialised the model at the beginning of each window before calculating the time-averaged population growth rate as percent change in coral cover per year. We stipulated that these windows must span at least 2 years in duration. Consequently, we identified three windows for each of the two reefs (Figure 7). For each recovery window, we reinitialised the model using the coral cover data from the nearest empirical data point. This ensured that both the model and empirical data growth rates were calculated from the same starting point.

We compared population growth rates as change in percentage coral cover over time for Moore and Thetford reefs for the: (i) Manta tow reef average with the average of model sites underlying the tow spatially, (ii) Manta tow average in each of the four sectors with the average of model sites underlying these sectors spatially (iii) Fixed photo transect average with the average of the three underlying or closest polygons.

The manta tow GPS track crossed areas that the geomorphic maps indicated were not the appropriate habitat for corals (i.e., not one of Reef Slope, Reef Crest, Outer Reef Flat or Sheltered Reef Slope) and were therefore not modelled explicitly by *C∼scape* in the case study. While we had GPS tracks of the manta tows, we did not have this information for each individual tow, and therefore could not remove the tows that were not over modelled reef areas for comparison to the model. Rather, we determined approximately how many site polygons would underlie the manta tow track if the geomorphic map has suggested that was coral-appropriate area. For Moore Reef this was eight sites in the back sector and for Thetford reef this was four sites in the back sector. For scenarios where we informed the model using the benthic habitat map, we specified that these sites have zero coral cover in all years.

## Acknowledgements

We acknowledge the Wulgurukaba, Bindal, Whadjuk Noongar, Turrbal and Jagera Peoples of the land on which the authors conducted the model development, analysis and writing for this study. We acknowledge the Gooreng Gooreng, Gurang, Bailai, Taribelang Bunda, Wunambal Gambera and Gunggandji peoples as the traditional owners of the Sea Country where the empirical data for this study was sourced. This work is part of The Reef Restoration and Adaptation Program which is funded by the partnership between the Australian Government’s Reef Trust and the Great Barrier Reef Foundation. The work presented here benefitted greatly from being part of the RRAP Modelling and Decision Support subprogram and from interacting with the EcoRRAP subprogram. We acknowledge the importance of the AIMS Long-Term monitoring program in allowing the model evaluation presented here. We greatly appreciative comments on manuscript drafts from Mariana Álvarez-Noriega and Mia Hoogenboom.

