## Supplementary Material for "Reproducing within-reef variability in coral dynamics with a metacommunity modelling framework"

Contents

Supplementary Information 1: Generating a spatially explicit seascape
Supplementary Information 2: Integral Projection Models
Supplementary Information 3: Acute disturbances
Supplementary Information 4: Moore Reef Cluster case study

### 1. Generating a spatially explicit seascape

#### 1.1. Delineating sites

As discussed in the main text, it was necessary to divide each reef(s) into units for model implementation. We used geomorphic maps from (Roelfsema *et al.* 2020). The geomorphic maps were created using a combination of machine learning and semi-automated expert-driven contextual editing (Lyons *et al.* 2020). A random forest classifier used expert-curated training data to predict geomorphic zones from a stack of Sentinel-2 satellite imagery (10 m pixels) and other physical attributes (depth, slope, wave environment parameters), the output of which was then contextually edited via an expert-driven ruleset.

Following the recommendations of Kennedy *et al.* (2021), we only considered the geomorphic zones that are expected to have predominantly hard substrate — Reef Slope, Reef Crest, Outer Reef Flat and Sheltered Reef Slope — to be areas where we would expect appropriate habitat and conditions for corals to grow (description in table S1, Roelfsema *et al.* 2021). While other zones may have patchy coral habitat, we considered this to be minor in terms of potential to contribute to the metapopulation, as it is likely that distinct species of corals occur in these more marginal habitats.

Figure S 1 illustrates the process of pixelating each geomorphic zone as detailed in the main text.

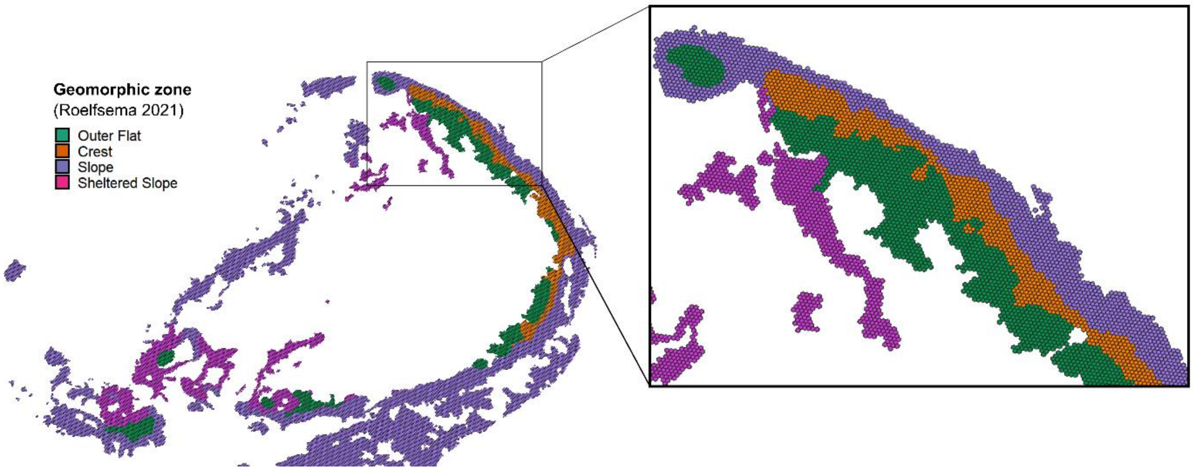


*Figure S 1. Moore Reef with the four selected geomorphic zones — Reef Slope, Reef Crest, Outer Reef Flat and Sheltered Reef Slope — pixelated into hexagons in preparation for clustering.*

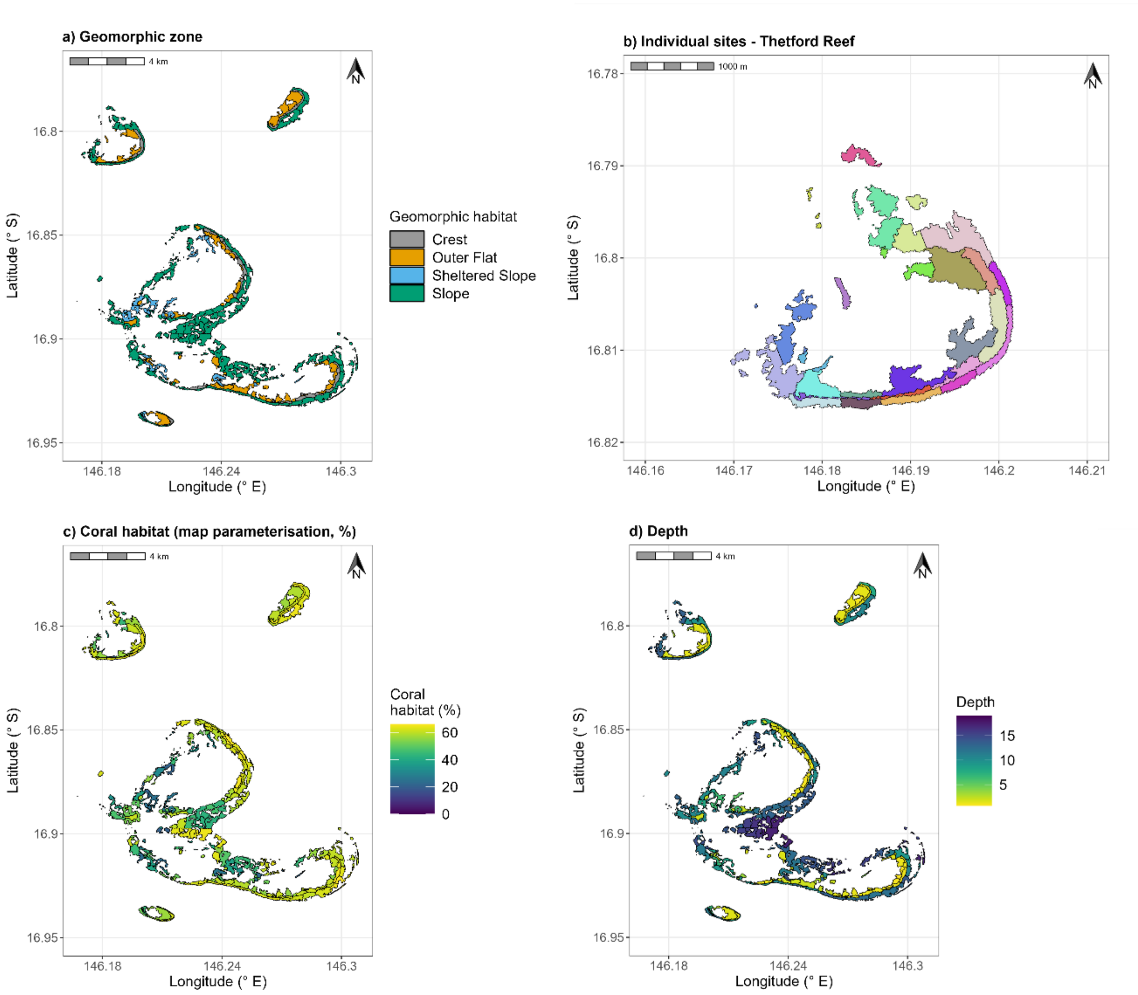


Figure S 2 shows the Moore Reef Cluster partitioned into sites for modelling.

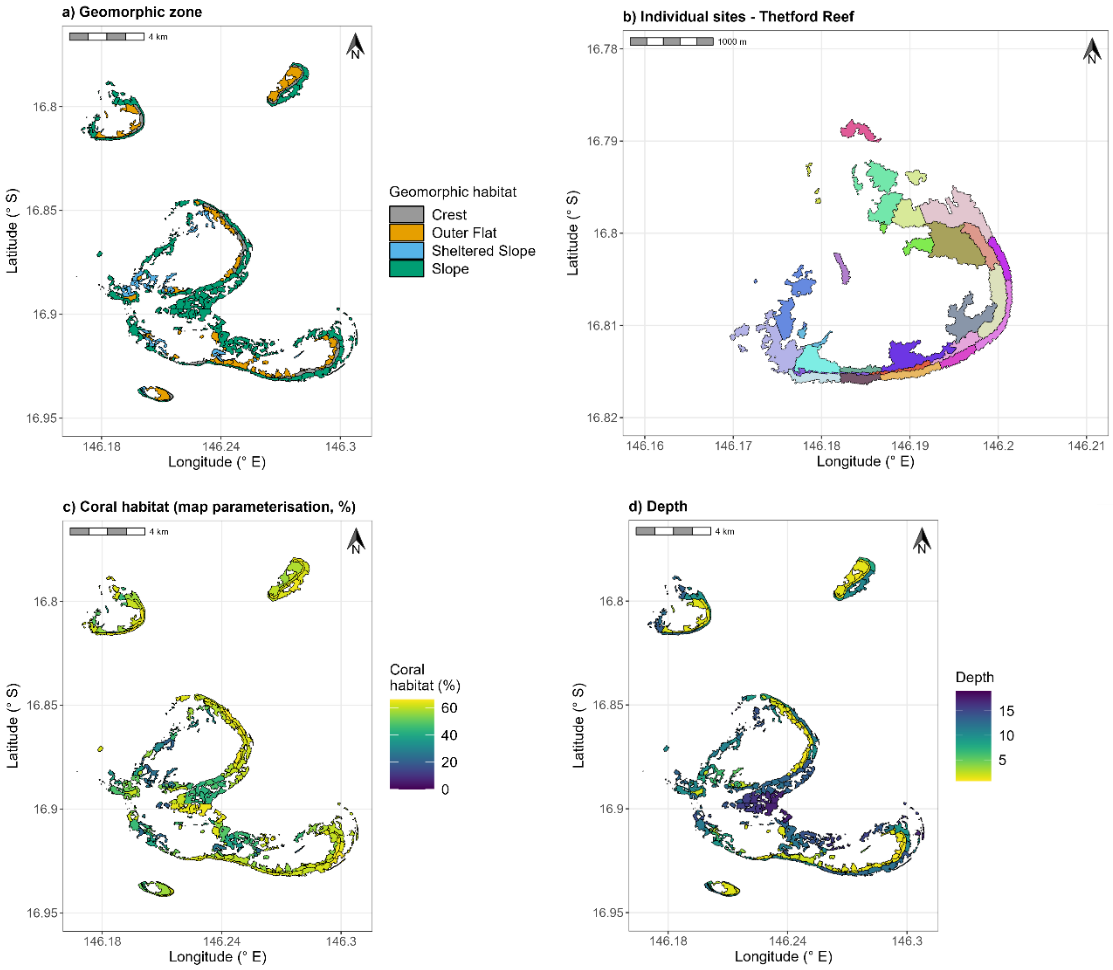


*Figure S 2. a) The Moore Reef Cluster partitioned into site polygons, coloured by geomorphic zone. b) Sites in Thetford Reef (north-west in Moore Reef Cluster) showing 27 individual sites. Each site is modelled discretely in C~scape and connected to the other polygons via the transport of coral larvae. Note that some polygons are 'multi-polygons', i.e., made up of >1 polygon, but still classified as one site for the modelling units. c) Sites coloured by their maximum coral habitat when parameterised by habitat maps. d) Mean depth (metres) of each polygon; depth influences the temperature stress experienced.*

| Reef | No. sites |
|---|---|
| Briggs | 7 |
| Elford | 95 |
| Milln | 15 |
| Moore | 69 |
| Thetford | 27 |
| **Total** | **213** |

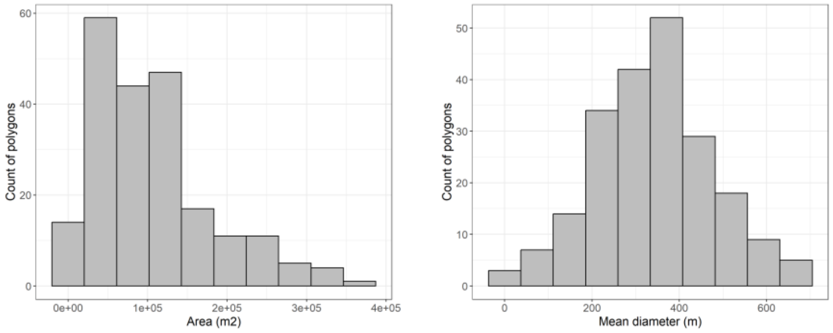


*Figure S 3. Summary of the number of site polygons in each reef and histograms showing the size distribution of the 213 modelled site polygons according to total area and diameter (when polygons are assumed to be circular). The mean diameter is 340 m2.*

#### 1.2. Parameterising site-specific coral habitat

The benthic habitat maps (Roelfsema *et al.* 2021) were used to calculate coral habitat. The benthic habitat maps were created using the same underlying framework and input data sets as the geomorphic zonation maps (Roelfsema *et al.* 2021). The key difference is the machine learning model was trained using point-based field observations and the contextual editing process was able to use the geomorphic map as an input layer.

Each site polygon was assigned a coral habitat value to represent the percentage of the total polygon area where coral could potentially grow. The benthic habitat maps from Roelfsema *et al.* (2021) is composed of 10x10m pixels classified as one of four categories to inform the calculation of coral habitat: Sand, Rubble, Rock, Coral/Algae (Figure S 4).

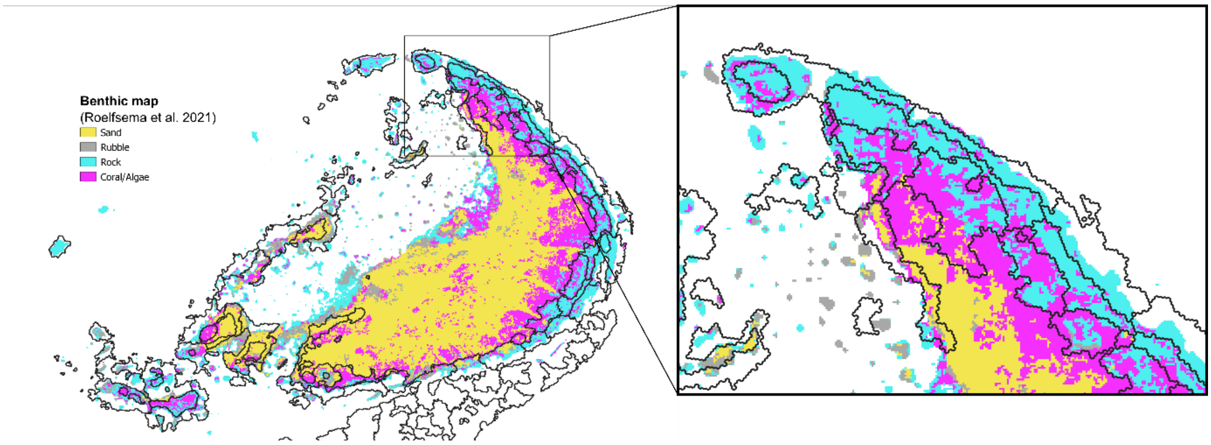


*Figure S 4. The benthic reef habitat from Roelfsema et al. 2021 showed for Moore Reef, is shown for Moore Reef, overlaid with the site polygons delineated from the geomorphic map in black.*

The benthic habitat map did not always cover the full extent of the geomorphic habitat map, meaning that there were pixels within the site polygons that had no benthic map information, i.e., NA values. These NA values were removed from the calculation of coral habitat. In cases where more than 95% of the pixels in any given polygon were NA, we instead took the value of coral habitat from the nearest neighbour polygon that had the same geomorphic classification.

All site polygons have an associated spatial area, $Area_i$, which varies depending on their size and shape. Importantly, this is distinct from the 'potential coral area' that is modelled, which can be calculated as the sum of each site area multiplied by its maximum coral habitat, $K_i$, as a proportion of its total area:

$Area_{reef} = \sum Area_i \times K_i$

#### 1.3. 3D reef area

Using the geomorphic maps to create the site polygons determines the total reef area which is modelled across the seascape. Assigning a coral habitat then sets an upper limit on the amount of coral area for each site polygon. We converted the 2D site polygon areas from the map to 3D surface area, using a "surface-to-horizontal-area ratio", derived from the slope value estimated for each pixel. Slope was estimated by using a local gradient method (3 × 3 window) from a bathymetric map. The slope-adjusted surface area, i.e., 3D surface area, was calculated using a trigonometric formula of 2D surface area and the slope where

$A_{3D} = A_{2D}/\cos(slope)$, where the slope is in radians.

The total 3D area of site polygons for Moore Reef Cluster was 22588866 $m^2$ = 226 hectares, but not all of this was available for coral, as governed by the coral habitat. If coral was at maximum coral habitat in all site polygons in the Moore Reef Cluster this would equate to 120.7 hectares of coral.

#### 1.4. Connectivity

The connectivity matrix captured variability in the number of larvae retained in a site, exported to other sites, and received from other sites (Figure S 5). Some reefs (e.g., Thetford Reef, northwest) were found to be strongly self-seeding, while other reefs (e.g., Thetford and Moore Reef) shared a lot of larvae (Figure S 5).

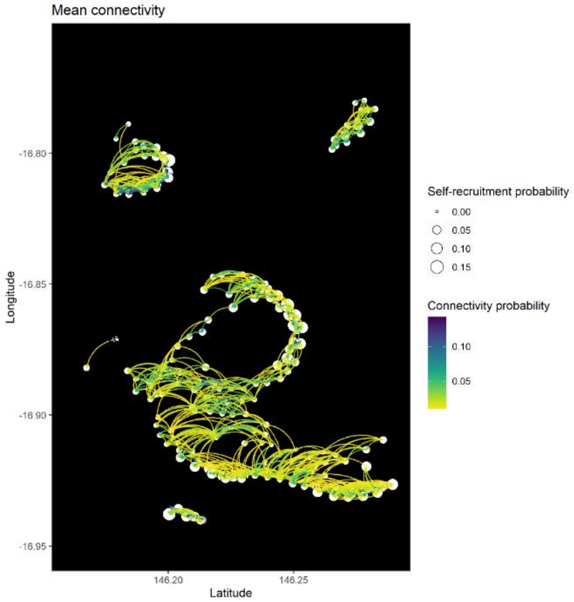


*Figure S 5. Visualisation of connectivity between the site polygons (represented by white dots). Colour of lines indicates proportion of larvae travelling between site polygons, while the size of white dots indicates proportion of larvae remaining within the source site polygon.*

### 2. Integral Projection Models

#### 2.1. Mathematical construction

An Integral Projection Model is a continuous-state, discrete-time model (Easterling et al. 2000) and is a form of an integrodifference equation (Neubert et al. 1995), where the spatial re-distribution is replaced by a size re-distribution. The Integral Projection Model consists of two components: the state variable and a transition matrix.

The state variable of the IPM describes the number of individuals $n(x,\ t)$ at year $t$ with state $x$. In this work, $x$ and $y$ represent the size as colony area of individual coral colonies as well as three additional discrete states: egg, larvae and settler such that:

$$n(x,\ t) = [n_{egg,t} \quad n_{larvae,t} \quad n_{settler,t} \quad n_{x\,1,t} \quad n_{x\,2,t} \quad n_{x\,3,t} \cdots \quad n_{x\,n-1,t} \quad n_{x\,n,t}]$$

This describes the abundance of coral eggs, larvae and settlers (the discrete states) and the size structure of the coral population at one point in time. The number of size classes, $n$, is user defined (see discussion of discretisation below).

The transition matrix, or kernal $k(y,x)$, is analogous to a projection matrix (e.g. Leslie matrix, (Hansen 1989)) in matrix population modelling (Caswell 2000). It represents all possible transitions from state $x$ (in year $t$) to state $y$ (in year $t+1$), integrated over all states of $x$ (Metcalf *et al.* 2013). The mathematical definition of the kernel is flexible but is here defined as the product of survival of individuals in state $x$ at time t, $s(x)$, and growth of individuals from state $x$ to state $y$, $g(x,y)$, plus the fecundity of individuals in state $x$ producing those in state $y$, $f(x,y)$, at time t+1.

$$k(y,x) = s(x)g(x,y) + f(x,y)$$

Survival, $s(x)$, is the probability that an individual of size $x$ at time t will be alive at time t+1. Growth, $g(y,\text{x})$, is the probability that an individual of size $x$ at time t will grow to an individual of size y at time t+1. This function considers cases of positive growth where $y > x$ and also negative growth, i.e., partial mortality of corals, where $y <$ x. The coral species we included generally did not fragment into multiple individual ramets, hence this was not captured in the demographic data and was not included in the models.

Similar to the growth and survival functions, fecundity, $f(x,y)$, is determined as a function of coral colony size. We assume that once a coral individual is large enough to reproduce, it reproduces once each year (once per model time step) until the individual dies.

The number of eggs that are produced by a coral colony is modelled as

$$f(x) = A\,\rho\,E\,m$$

Where $A$ is the colony surface area in $\text{cm}^2$, $\rho$ is the density of coral polyps ($\text{cm}^{-2}$) (Doropolous et al. 2020), $E$ is the number of eggs per polyp (Pratchett et al. 2019; Doropolous et al. 2020), and $m$ is the proportion of these polyps that are mature (Alvarez-Noriega et al. 2016, Doropoulos et al 2020).

The kernel $k(y,x)$ can be used to predict the number of individuals $n(y,\ t+1)$ at year $t+1$ with a given state $y$ as a function of the number of individuals $n(x,\ t)$ at year $t$ with state $x$.

$$n(y,\ t+1) = \int_L^U k(y,x)n(x,t)dx$$

Discretisation is necessary to facilitate the numerical evaluation of the integral. A meshpoint, $x$, serves as the mid-point for each bin in the discretisation, representing a size class within the lower (L) and upper (U) size of the coral. We used 1 cm diameter for the lower limit for the continuous state (capturing corals smaller than this in the 'settler' discrete state). The upper diameter is set at 90% of the maximum size observed in the data at the beginning of the sampling period to avoid statistical predictions at the limits with few samples. There is a trade-off between using a high number of meshpoints ($m$), which should give greater accuracy, and the computational cost of a high number of meshpoints. In the present study we used 100 meshpoints, which were evenly distributed on the log-scale of coral colony surface area.

The width of a given meshpoint can be calculated as $\Delta x = \frac{(U-L)}{m}$. Combining this with the midpoint rule (Ellner & Rees, 2006) the numerical evaluation results in

$$n(y, t+1) = \Delta x \sum_{i=1}^{m} k(\mathrm{y}, \mathrm{x},) n(\mathrm{x}, t)$$

Which gives a matrix, *m* by *m,* here 100 by 100, that represents the continuous state transition matrix (Metcalf *et al.* 2013).

The discrete transitions must also be parameterised, i.e., the transition probability from eggs to larvae, larvae to settlers, and settlers into the continuous state (see Figure S 6).

#### 2.2. The coral life cycle

An integral projection model needs to capture the core processes in an organism's life cycle over a temporal period, here annual, such that changes in the abundance and size, and mortality and recruitment to a population can be synthesised into a transition probability matrix. Figure S 6 gives a schematic summary of the coral life cycle as modelled in this study.

Growth, survival, and fecundity could be modeled as a function of coral size as a continuous variable, but we also required several discrete transitions. Therefore, we included discrete states for coral egg, larvae, and settler to close the life cycle and complete the integral projection matrix (Merow *et al.* 2014). As in many demographic studies, the early life history is relatively poorly understood (Elderd & Miller 2016), despite the importance of processes such as fertilization, settlement, and survival of recruits in determining population success (Hughes & Jackson 1985; Babcock 1991; Doropoulos *et al.* 2015). We parameterised these transitions with information from the literature (Table S 1).

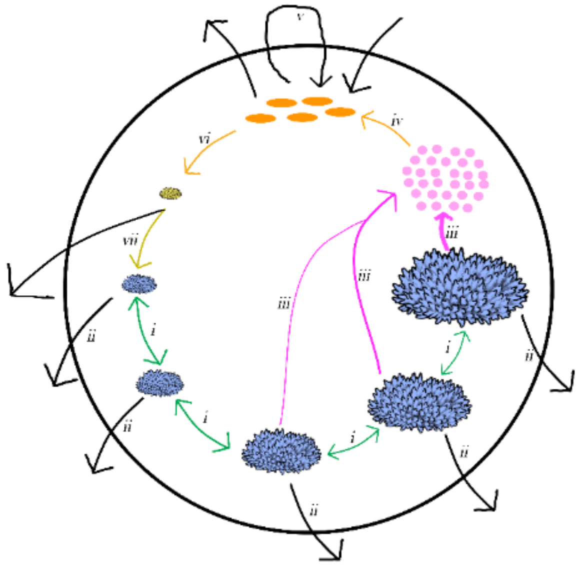

*Figure S 6. A schematic representation of the coral life cycle and the main modelled processes. Different sized coral colonies are represented in the lower hemisphere of the circular life cycle, as well as an egg, larvae and settler state in the upper hemisphere. In any given timestep a coral may die (ii, survival), or else may transition to a different size (i, growth), either growing, shrinking (partial mortality) or staying the same size, with probabilities determined by statistical Bayesian regressions of growth and survival. Mature colonies may reproduce once in a timestep (iii), releasing gametes into the water column which may fertilise to become larvae (iv). Transport of these larvae is then simulated separately via connectivity modelling, before these larvae may settle within a site to become a settler (vi), before transitioning into the continuous state one year later (vii). The approach for modelling these processes and the data used are detailed in Table S1.*

#### 2.3. Vital rate regressions: growth and survival

To construct the integral projection matrices, growth, survival and fecundity functions are obtained from statistical regressions built using empirical data from monitoring coral colonies over annual timesteps (growth and survival functions) and surveying the number of eggs and polyps in coral colonies (fecundity function).

We fit the models for each of growth, survival, and fecundity as a function of coral size in R using the brms package (Bürkner 2017). We used multilevel models to predict the three continuous vital rates as a function of coral colony area ($cm^2$). We followed the approach of Elderd and Miller (2016) and Kayal *et al.* (2018) to fit the models using a Bayesian framework.

Colony growth was modelled by predicting coral colony surface area at time t+1 as a function of coral colony surface area at time t, on a natural log-log scale, assuming a Gaussian distribution with identity link. We included a random effect for colony ID, to account for some colonies being measured over multiple years. We included a fixed effect for dataset for the corymbose *Acropora* model. *Goniastrea* data was only available from Scott Reef and so we did not require a fixed effect for this model.

Survival models were fitted using a Bernoulli distribution (and a logit link) with binary output of alive or dead at time t+1 as a function of colony area (on the natural log scale) at time t. Fixed and random effects were incorporated as for the growth models.

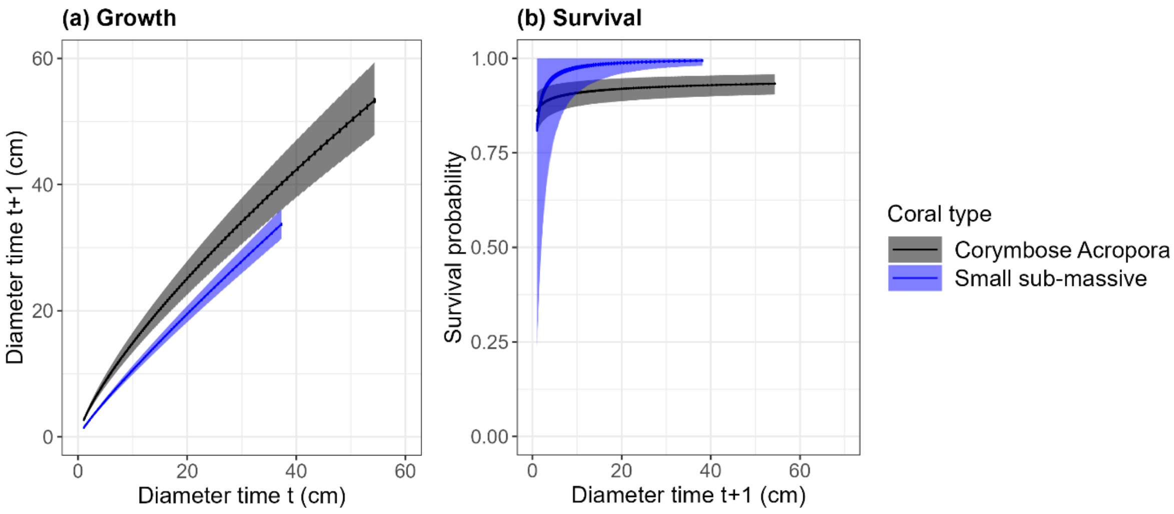


*Figure S 7. Predictions from a) growth and b) survival statistical regressions. a) Predicted diameter at time t+1 as a function of diameter at time t and b) predicted probability of surviving the annual period as a function of coral colony diameter at time t. Predictions are shown in grey for corymbose Acropora and blue for the small-massive Goniastrea coral types. Solid line shows mean and the ribbon shows the 95% Credible Interval.*

#### 2.4. Vital rate regressions: fecundity

Fecundity (the number of eggs produced by a coral colony, *E*) was modelled as a function of colony area. We used a Hurdle-Poisson model (with a log link) which could account for small colonies being non-reproductive. Hurdle models are commonly used to model zero-inflated ecological data (Balderama et al. 2016; Brown et al. 2016; Cunningham et al. 2018) and consist of two components. The first component is the zero-inflated model, which describes the probability of coral being reproductive or not. The second component is the conditional model, which uses a zero-truncated error distribution to describe the relationship between model predictors and non-zero egg count data.

We separately fitted a model the number of polyps per $cm^2$ colony surface area, $\rho$ as function of coral size in $cm^2$ (log-transformed) with a gamma distribution and a log-link, to account for larger colonies potentially having more or less polyps per surface area.

Multiplying predictions from these models was used to estimate the number of eggs as a function of colony size (Figure S 8).

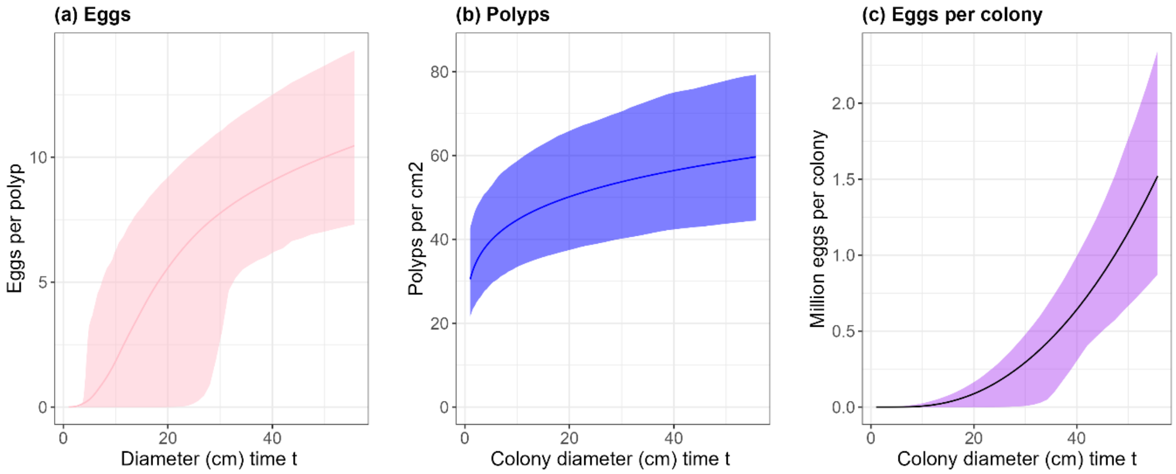


*Figure S 8.a) Eggs per polyp as a function of coral colony size. Notably, at approximately 10 cm diameter colonies start to become reproductive. B) polyps per cm2 of coral area as a function of colony size. c) multiplying the predictions from the regressions in a) and b) gives a prediction of the number of eggs produced per colony. Solid line shows the mean, and the ribbon shows the 95% Credible Interval. These results were used for both corymbose Acropora and for the small-massive Goniastrea coral types.*

#### 2.5. Capturing uncertainty from the Bayesian regressions

Predicting from the regressions allows the creation of integral projection matrices. We sampled growth, survival and fecundity predictions from the joint posterior distributions of the model 100 times to maintain a measure of uncertainty in vital, i.e.:

$$k(y,x)_i = s(x)_i g(x,y)_i + f(x,y)_i$$

where $k(y,x)_i$ is a kernel or integral projection matrix, $s(x)$ is survival of individuals in state $x$ from time t to t+1, $g(x,y)$ is the growth of individuals from state $x$ to state $y$, $f(x,y)$ is the fecundity of individuals in state $x$ producing those in state $y$ at t+1, and $i$ is a posterior sample.

#### 2.6. Discrete transitions

Probabilities for egg fertilisation to larvae, and larvae settlement probability were informed from the literature (see Table S 1).

The transition from settler to the continuous state of the IPM required data on what size the settler corals were likely to be 1-year following settlement, as this would define their size, $n(y,\ t+1)$. To parameterise this we used data from Cruz and Harrison (2017), Dela Cruz and Harrison (2020) and Harrison *et al.* (2021) who measured the survival and growth of corals that settled on tiles and natural substrates over ~3 years.

To parameterise the size at which corals would transition from settler into the continuous state of the integral projection matrix we generated their randomly from a normal distribution with mean value 2.56 cm for acroporids and 1.41 cm for non-acroporid species according to Cruz and Harrison (2017), Dela Cruz and Harrison (2020) and Harrison *et al.* (2021). We used a standard error of 0.1 times the mean diameter.

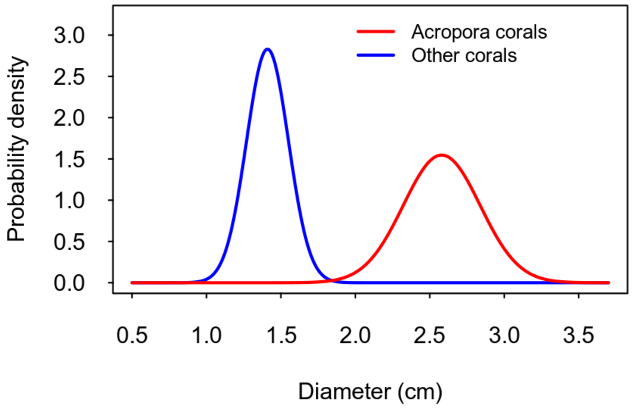


*Figure S 9. Probability density plot showing the distribution which was sampled to determine the probability of transitioning to each size class in the integral projection matrix 1 year after settling.*

*Table S 1. Summary of the vital state transitions modelled in the Integral Projection Models to capture the life cycle shown in Figure S 6. The data and approach used to model or parameterise the transition is detailed and references are provided.*

| Vital rate transition | Process | Data used to | Parameters/ data corymbose *Acropora* | Parameters/ data small sub-massive *Goniastrea* | Reference(s) |
|---|---|---|---|---|---|
| i | Growth/ Shrinkage (partial mortality) | Fit regressions for $\text{size}_{t+1} \sim \text{size}_t$ | Growth data encompassed a set of broadly comparable *Acropora* species across three datasets. Data from 2329 coral colonies from Scott Reef[a] (A.millepora), 2045 from Moorea[b] (*Acropora hyacinthus, A. globiceps, A. retusa*, and *A. fragilis*), 202 from Heron Island, GBR[c] (colonies <5cm diameter at time t: *A. hyacinthus, A. nasuta, A. humilis, and A.* spp). Site-year combinations that had known acute disturbances in the Scott Reef dataset were filtered out a priori. | Data for *Goniastrea* was sourced only from the Scott Reef dataset only and included *G. retiformis* and *G. edwardsii*. Data was obtained from 1054 coral colonies from Scott Reef[a]. Site-year combinations that had known acute disturbances in the Scott Reef dataset were filtered out a priori. | [a] Gilmour et al. (2013) [b] Kayal *et al.* (2018) [c] Doropoulos et al. (2015) |
| ii | Survival | Fit regressions for $\text{survivalstatus}_{t+1} \sim \text{size}_t$ | As above. | As above. | As above. |
| iii | Fecundity: eggs per polyp | Fit regressions for number of eggs per polyp as function of colony size $F_{oocyte_{t+1}} \sim size_t$ | Data was taken from on average 5 polyps from 3 separate branches from 120 A. millepora coral colonies total from Scott Reef. | As for Acropora. | Gilmour et al. (2013) Foster and Gilmour (2020) |
| | Fecundity: polyps per cm² coral area | Fit regressions for number of polyps per colony surface area as function of colony size $F_{polyp_{t+1}} \sim size_t$ | Data was taken from 120 A. millepora coral colonies total from Scott Reef. | As for Acropora. | Gilmour et al. (2013) Foster and Gilmour (2020) |
| | Fecundity: colony | $F_{colony_{t+1}} \sim F_{oocyte}(size_t) \times F_{polyp}(size_t)$ | As above. | As above. | As above. |
| | Polyp sexual maturity | Parameterise the proportion of mature polyps per colony. Multiplied by $F_{colony_{t+1}}$ to account for unmature polyps within a colony. | 40% of polyps assumed to contain fertile eggs. | As for Acropora. | Doropoulos *et al.* (2019) Álvarez-Noriega *et al.* (2016) |
| iv | Fertilisation to larvae | Parameterise fertilisation probability: multiplied by $F_{colony_{t+1}}$ to determine number of larvae. | 55% of spawned eggs assumed to get fertilised in the water column. | As for Acropora. | Oliver and Babcock 1992 . |
| | Early mortality prior to dispersal | Parameterise larvae mortality: multiplied by $F_{colony_{t+1}}$ to determine number of larvae. | 50% of larvae assumed to die in first three days during dispersal, before becoming competent to settle (e.g. predators, natural mortality) | | Graham, E. M., Baird, A. H., & Connolly, S. R. (2008). |
| v | Larvae transport, export and import | Connectivity model conducted separately to create connectivity matrix describing probability of moving between each and every site population polygon. | See main text | See main text | |
| vi | Settlement | Parameterise the probability that a larvae will settle if it is over reef substrate within its competency window. | From the lower and upper limits of larvae settlement probabilities[d,e] we took the mean probability of settlement for Acropora to be 2.1% ((0.05-0.008)/2): 2.1% (Acropora) | The settlement probability of Goniastrea was assumed to be 60% that of Acropora 1.26% (Goniastrea)[f] | [d] Edwards et al. 2015 [e] De La Cruz and [f] Harrison (2017), Wallace (1985) |

| | | | | | |
|---|---|---|---|---|---|
| vii | Growth to 1-year old coral | Parameterise the probability of transitioning from the discrete ‘settler’ state to the continuous state. | 1.315% | 1.315% | Doropoulos *et al.* (2019) |
| vii | Size at 1-year old | Used to specify the size of corals 1 year after settlement. It is at this point that they transition into the continuous state of the IPM and must be assigned a size. | Sampled from a normal distribution with mean 2.56cm and standard deviation 0.26cm. | Sampled from a normal distribution with mean 1.41cm diameter and standard deviation 0.14cm | Cruz and Harrison (2017) Dela Cruz and Harrison (2020) Harrison *et al.* (2021) |

### 3. Coral population mortality agents

#### 3.1. Temperature stress

***Temperature stress spatial variability***

For the purposes here, the eReefs RECOM model (Herzfeld 2009; Steven *et al.* 2019) nested within GBR1 (Skerratt *et al.* 2023) was used to simulate transport and dispersal of larvae with a ~250 m spatial resolution. RECOM was used to implement a hydrodynamic model in SHOC (Herzfeld 2009) with a grid resolution of approximately 250 m within the 1 km resolution version 2.0 GBR-scale hydrodynamic model. SHOC can be used to calculate two- or three-dimensional DHWs given a reference climatology. For this project, we used the 4 a.m. DHW product using the STTAARS regional climatology (Wijffels *et al.* 2018).

While this product is not entirely consistent with the NOAA Coral Reef Watch product, and an improved approach is needed in future work, it was used only to estimate the relative differences between grid-cells on the local scale, for local-scale adjustment of regional DHW projections.

We calculated the proportional residuals (R) between the mean DHW value of a reef and all sites within the reed for the three marine heatwave years that were modelled using RECOM:

$$R_{i,y} = \frac{DHW_i - \frac{\sum DHW_i}{n}}{DHW_i}$$

where $i$=1 to $n$ for each of the 213 sites in the Moore Reef Cluster and $y$ is each of the marine heatwave years (2016, 2017, 2020).

Next, the mean, $mean(R_i)$, and standard deviation, $std(R_i)$, were calculated for each site across the three years to allow the fitting of a normal distribution.

We could then assign $DHW_i$ to each site for each time, $t$, when we had a hindcast value for the DHW experienced at the reef level ($DHW_{reef,t}$) by using the normal distribution of the residuals:

$$DHW_{i,t} = DHW_{reef,t} + DHW_{reef,t} * rnorm(mean(R_i), std(R_i))$$

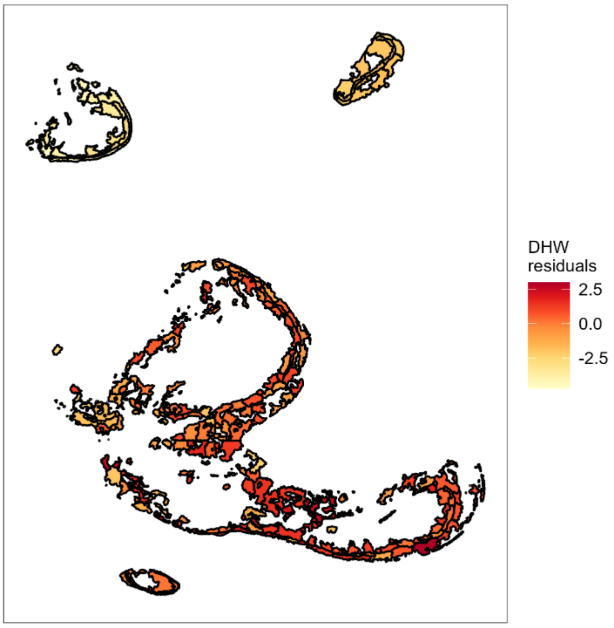


*Figure S 10. The mean Degree Heating Week residuals across the three marine heatwave years examined showing how spatial variation in mean heat exposure is implemented in the C~scape framework.*

***Temperature stress mortality in C~scape***

The probability of coral mortality in any given year was estimated as a function of DHW following the model developed in Bozec *et al.* (2022) based on observations of coral mortality during the 2016 bleaching event (Hughes *et al.* 2018). The data available from (Hughes *et al.* 2018) data could be divided into two types: (i) initial bleaching mortality recorded at the time of peak temperature stress and (ii) long-term mortality observed after 6 months. Due to the yearly time-step in C~scape, long-term mortality was the most appropriate metric to use, but was in the form of a change to coral cover. Initial bleaching data was at the colony level, which was more appropriate given the IPM engine of C~scape. Therefore, we used both data types following the approach of Bozec *et al.* (2022) and calculated bleaching mortality in two main steps.

First, mortality at the peak of a bleaching event was calculated using a model of initial mortality as a function of DHW, fitted to the data from figure 2a Hughes *et al.* (2018) and based on the equations developed for ReefMod-GBR (Bozec *et al.* 2022).

Two additional coefficients were added to the model of initial mortality such that

$$m_{init} = w * s * e^{(0.17+0.35*DHW)-1}/100$$

with $w$ being the depth coefficient and $s$ being the bleaching sensitivity of a species.

The bleaching sensitivity coefficient was included because the data from figure 2a, Hughes *et al.* (2018) was from multiple different coral types and we needed to apply it more specifically to the coral types modelled in the present study so we followed Bozec *et al.* (2022) approach here.

Here, corymbose *Acropora* is given a bleaching sensitivity of 1.5, while small massive *Goniastrea* is 0.25.

The depth coefficient was included because data in figure 2a, Hughes *et al.* (2018) was collected at roughly 2m depth and is not representative of all depths.

Another piece of information from Baird *et al.* (2018), showing the relationship of bleaching with depth (fig 2, (Baird et al. 2018), was included to obtain depth coefficient $w$. We fit an exponential model to the data from fig 2, Baird *et al.* (2018), to get a formula to calculate the depth coefficient $w$ using the depth of each site.

$$w_{site} = e^{-0.07551(sitedepth-2)}$$

The sensitivity for each coral type was taken from table S1, Bozec *et al.* (2022): the scaling was 1.4 for corymbose / small branching Acroporids (here used for corymbose *Acropora*) and 0.25 for small sub-massive corals (here used for small sub-massive *Goniastrea*).

The initial mortality of each coral at each site is therefore calculated as

$$m_{init_{ft\_site}} = w_{site} * s_{ft} * ((e^{0.17+0.35*DHW_{site}}) - 1)/100$$

$m_{init_{ft\_site}}$ was capped at 1.

Annual mortality ($M$) was then calculated based on data from figure 2c Hughes *et al.* (2018) and work by Bozec *et al.* (2022) who calibrated the long-term bleaching mortality to the initial bleaching mortality by combining information from figure 2a and 2c in Hughes *et al.* (2018).

$$M_{ft\_site} = 1 - (1 - m_{init_{ft\_site}})^6$$

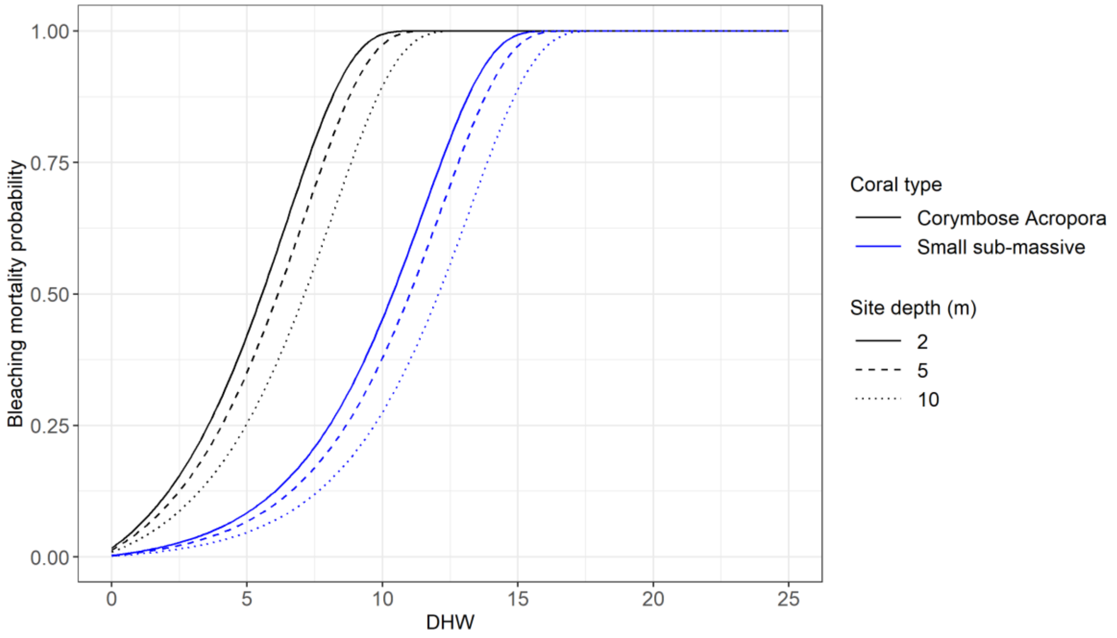

#### 3.2. Cyclones as mortality agent

As in Bozec *et al.* (2022), past exposure to cyclones was derived from sea-state predictions of wave height (Puotinen et al. 2016). The potential for coral-damaging sea state (wave height >4 m) was determined using a map of wind speed every hour within 4-km pixels over the GBR for cyclones between 2008 and 2020. Any reef containing a combination of wind speed and duration capable of generating 4-m waves, assuming sufficient fetch, was scored as positive for potential coral-damaging sea state in the respective year. Where damaging waves were predicted, an estimate of cyclone category was deduced from the distance to the cyclone track extracted from the BoM historical database (http://www.bom.gov.au/cyclone/tropical-cyclone-knowledge-centre/understanding/tc-info/)

*Table S 2. Cyclone categories and their associated windspeed, according to the Bureau of Meteorology, used to make a conversion from cyclone category to windspeed.*

| Cyclone category | Windspeed |
|---|---|
| 1 | 24.5 m/s |
| 2 | 32.5 m/s |
| 3 | 44.2 m/s |
| 4 | 55.3 m/s |
| 5 | 65 m/s |

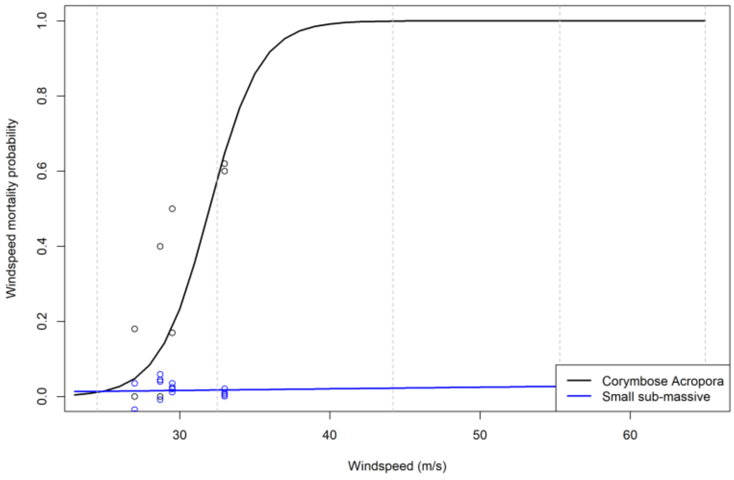


*Figure S 11. Windspeed and the predicted associated probability of coral mortality for the two coral types included in this study: corymbose Acropora and small sub-massive Goniastrea. Dashed vertical lines indicate cyclone categories 1-5 from left to right.*

#### 3.3. Crown of thorns starfish as a mortality agent

We calculated the feeding on corals by COTS and the resulting coral mortality following the approach in Bozec *et al.* (2022). This approach was informed by published rates of consumption based on the size of COTS in Keesing and Lucas (1992), converted to the eight age classes using information in Engelhardt *et al.* (2001).

To include COTS consumption selectivity between coral types, information from De'ath and Moran (1998) is utilised. According to this data, *Acropora* is preferred over *Goniastrea* at an odd ratio of 14:4.3.

### 4. Moore Reef Cluster case study

#### 4.1. Simulation details

*Table S 3. Details of model simulations. All simulations in the table were repeated twice, once for each of the two parameterisation of coral habitat. Simulations were started in different years reflecting the start of each recovery window (Figure 7) examined to obtain the time-averaged annual change in coral cover.*

| Dataset for validation and initialisation | Year initialised | Location of initialisation data | Recovery windows examined for reef and reef sector |
|---|---|---|---|
| Fixed-position photo-transects | 2008 * | Moore & Thetford slope | 2008-10 Moore<br>2008-10 Thetford |
| Fixed-position photo-transects | 2012 | Moore & Thetford slope | 2012-16 Moore<br>2012-16 Thetford |
| Fixed-position photo-transects | 2018 | Moore & Thetford slope | 2018-20 Moore<br>2018-21 Thetford |
| Manta tow | 2008 | Thetford | 2008-10 Thetford<br>2008-10 Thetford Front<br>2008-10 Thetford Back<br>2008-10 Thetford Flank1<br>2008-10 Thetford Flank2 |
| Manta tow | 2012 | Moore & Thetford | 2012-16 Thetford<br>2012-16 Thetford Front<br>2012-16 Thetford Back<br>2012-16 Thetford Flank1<br>2012-16 Thetford Flank2<br>2012-16 Moore<br>2012-16 Moore Front<br>2012-16 Moore Back<br>2012-16 Moore Flank1<br>2012-16 Moore Flank2 |
| Manta tow | 2018 | Moore & Thetford | 2018-22 Thetford<br>2018-22 Thetford Front<br>2018-22 Thetford Back<br>2018-22 Thetford Flank1<br>2018-22 Thetford Flank2<br>2018-21 Moore<br>2018-21 Moore Front<br>2018-21 Moore Back<br>2018-21 Moore Flank1<br>2018-21 Moore Flank2 |

#### 4.2. Initialisation

As detailed in main text, simulations were initialised based on coral cover recorded in the AIMS LTMP observations from the starting year of the hindcast (2008 for the full trajectory, or the start of each 'recovery window' for the population growth analysis). For the manta tow evaluation, coral cover was based on the manta tow observations, while the initial composition of the corymbose Acropora to the small sub-massive Goniastrea group was based on the ratio observed in the phototransect dataset at the initialisation year. Any site polygons directly underlying the sites of the phototransects or the manta tows were assigned the values matching to the phototransect/tows. Initialisation for sites not underlying a phototransect or tow, was extrapolated based on the coral habitat assigned to a given site and the ratio of coral cover to the maximum coral habitat. The coral habitat ratio (KR) was calculated for each site matched with a phototransect or tow in the previous step, using the total coral cover of the site and its coral habitat

$$KR_i = \frac{T_{cover\_site_i}}{K_i}\ .$$

This allowed us to form a normal distribution N(μ=mean(KR), σ =sd(KR)) for each reef within the cluster. Finally, to extrapolate initial total coral cover for each site, we draw a sample KR from this distribution and calculate the coral cover:

$$T_i = KR_i \times K_i$$
